# Allostery in Proteins as Point-to-Point Telecommunication in a Network: Frequency Decomposed Signal-to-Noise Ratio and Channel Capacity Analysis

**DOI:** 10.1101/2021.02.26.433010

**Authors:** Yasemin Bozkurt Varolgüneş, Joseph F. Rudzinski, Alper Demir

## Abstract

Allostery in proteins is a phenomenon in which the binding of a ligand induces alterations in the activity of remote functional sites. This can be conceptually viewed as *point-to-point telecommunication in a networked communication medium*, where a signal (ligand) arriving at the input (binding site) propagates through the network (interconnected and interacting atoms) to reach the output (remote functional site). The reliable transmission of the signal to distal points occurs despite all the disturbances (noise) affecting the protein. Based on this point of view, we propose a computational frequency-domain framework to characterize the displacements and the fluctuations in a region within the protein, originating from the ligand excitation at the binding site and noise, respectively. We characterize the displacements in the presence of the ligand, and the fluctuations in its absence. In the former case, the effect of the ligand is modeled as an external dynamic oscillatory force excitation, whereas in the latter, the sole source of fluctuations is the noise arising from the interactions with the surrounding medium that is further shaped by the internal protein network dynamics. We introduce the excitation frequency as a key factor in a *Signal-to-Noise ratio (SNR)* based analysis, where SNR is defined as the ratio of the displacements stemming from only the ligand to the fluctuations due to noise alone. We then employ an information-theoretic (communication) channel capacity analysis that extends the SNR based characterization by providing a route for discovering new allosteric regions. We demonstrate the potential utility of the proposed methods for the representative PDZ3 protein.

## 1 Introduction

The coupled dynamics of biological molecules underlie essential functions in cellular processes, including protein regulation and cellular signaling. Allostery—the transmission of the effect of ligand binding to another (often) distal site of a protein—plays an important role in many of these functions and may also be leveraged for novel drug delivery applications [1, 2]. There has been sustained activity in both computational and experimental allostery research ever since Monod and Jacob introduced the concept [3]. Nevertheless, a general mechanistic understanding of allosteric processes still remains elusive [4].

Since its introduction, the definition of allostery has evolved over time. In the earlier structure-centric models, the ligand induced change in the binding affinity at a distal site was thought to be accompanied by significant conformational changes. Two historically dominant models for allosteric mechanisms are the Monod–Wyman–Changeux (MWC) [5] and the Koshland–Nemethy–Filmer (KNF) models [6]. Both models assume that there is an equilibrium between two pre-existing conformational configurations, corresponding to the active and inactive states, but differ in the treatment of intermediate states. In the MWC model, all subunits simultaneously transition from the inactive to the active configuration upon ligand binding, i.e., they undergo a *concerted* motion. On the contrary, in the KNF model, the subunits transition, one at a time, in a *sequential* manner, resulting in transition states. Despite their initial success, these structure-based models are now considered insufficient for explaining the behavior of some allosteric proteins, such as the PDZ domains. As a result, a variety of extended models have been proposed. For instance, in the population-shift model, instead of only two states, proteins are assumed to exist in an ensemble of conformations. Subsequent to ligand binding, the ensemble undergoes a population shift towards a state that is favored by the ligand [7]. A more recent view called *dynamic allostery*, that was introduced by Cooper and Dryden, led to a paradigm shift in the notion of allostery by showing that, even in the absence of an observable conformational change in the mean structure, some proteins can exhibit allosteric behavior [8]. This result suggests that all proteins may be considered as intrinsically allosteric [9]. Cooper and Dryden further demonstrated via statistical thermodynamics based analysis that cooperative interaction free energies could deviate on the order a few kJ/mol upon ligand binding as a result of the changes in the frequency and amplitude of thermal fluctuations, with only a subtle change in the mean structure [8]. This result suggests that a frequency domain analysis of the fluctuations may provide quantitative insight into the detailed mechanisms of action-at-a-distance phenomena in allostery.

Motivated by the work of Cooper and Dryden, the present study proposes a set of novel computational methods for investigating allosteric processes by examining the effect of perturbations, due to both ligand binding and noise, over a range of relevant frequencies. To facilitate the scanning of perturbation frequencies, we represent the protein as a mass-spring network, as a graph of nodes connected by edges. The nodes of the network correspond to the constituting atoms, or coarse-grained beads (defined as a collection of atoms), and the edges represent the interactions between them. Noise forces are applied to all nodes, to model the thermal fluctuations that originate from the protein-solvent interactions. Similarly, the ligand-protein interaction is represented by external forces, but they only act on the nodes that are in close contact with the binding site. The response at a (distal) node selected as the output is then determined by the noise forces acting on all of the nodes that are further shaped by the network dynamics, as well as the external forces representing the ligand. When “signal transmission” from a specific input to multiple output sites is considered, signal attenuation, and transmission reliability that is also affected by noise, may vary a great deal over the output sites. Accordingly, the emergence of allosteric response due to ligand binding may be attributed to the *Signal-to-Noise ratio* (SNR) characteristics measured at the output site. We put forward that, in order for an allosteric response to occur at a certain output site, the associated SNR value must be relatively high.

Instead of treating ligand binding as a static structural event, we capture the dynamic nature of the ligand-protein interactions via modeling the external forces as dynamic oscillatory excitations, as in our recent previous work on frequency domain perturbation-response characterization for proteins [10], and as in related work [11]. While the excitation frequency is swept over a relevant frequency range, the frequency dependence of SNR is fully characterized. Instead of considering only the low-frequency (global) modes of motion, we argue that a full spectrum analysis is essential, especially in cases where SNR does not follow a monotonic trend as a function of frequency. In particular, noticeably higher SNR within a particular frequency band may be the key in identifying dynamic allosteric response.

Akin to network representation of proteins, frequency decomposed SNR based analysis further enables the conceptualization of allostery as a *point-to-point channel* in a multi-channel, networked, noisy communication medium, pointing to the direct link between SNR and *channel capacity* that was established by Shannon. In the communications and information theory framework, the ligand serves as a transmitter (source of information), while the whole protein acts as the noisy medium of communication, i.e., the channel, and the region where the response is probed can be regarded as the receiver (of information). As in all communication systems subject to noise, there is an upper limit for the error-free (reliable, effective) information transmission rate from the transmitter to the receiver, defined as the channel capacity. Based on the Shannon-Hartley theorem, the channel capacity is determined by the integral of SNR over the relevant frequency band [12]. As an extension to frequency decomposed SNR analysis, we propose to utilize channel capacity as an aggregate (over frequency) and interpretable measure of robustness of the communication between the ligand binding site and the distal regions of the protein in the presence of noise.

Previous efforts attempting to decipher allostery have mostly concentrated on the identification of *critical* residues, i.e., residues whose activities are most affected by the binding event. We propose to add another layer to this analysis by anatomizing each residue’s frequency decomposed SNR profile individually. Furthermore, we propose two different methods for characterizing the channel capacity, namely, the *per residue scan* and the *binding pocket excitation* schemes. These methods differ in terms of the number of nodes upon which the external force is applied, as well as with respect to the excitation characteristics. Each analysis aims to detect distinct features. In the per residue scan, while the external force is applied to each (input) node, one at a time, the channel capacity values are calculated at all of the remaining nodes as candidate outputs. Through this analysis, the input-output residue pairs that have the potential to exhibit allosteric coupling can be identified without relying on any prior site-specific binding information.

Hence, this analysis can be used to discover novel targets for ligand binding. In the binding pocket excitation scheme, the external force is applied specifically to the residues that are known to be located in the binding pocket of a specific ligand, while the channel capacities for the rest of the protein residues are calculated. The aim here is to identify the residues that are most likely to allosterically interact with an already known binding event.

The utility of the proposed methods is presented for a representative single-domain allosteric protein that is known to display dynamic allostery, namely the third PDZ domain of the PSD-95 protein. For an overview of previous approaches applied to PDZ3, the reader is referred to [2]. This study aims to contribute to this growing area of research through novel frequency-decomposed SNR and channel capacity based analysis techniques, which provide complementary insight with respect to previous methods. The proposed techniques have the potential to offer unique perspectives in probing the responses to external excitations while also taking the noise characteristics into account.

The paper is organized as follows: Section 2 describes the theoretical foundations, and Section 3 presents the proposed and employed methods. Section 4 shows the results for the investigation of allostery in PDZ3. Section 5 presents our conclusions. Supplementary material and details are provided in the Supplementary Information.

## 2 Theory

We describe the Signal-to-Noise ratio (SNR) and the channel capacity formulations in a step-by-step manner in the following subsections. Section 2.1 presents the Langevin formulation for the network dynamics driven by thermal noise and external forces. The stochastic properties of the thermal fluctuations are also discussed. Then, Section 2.2 introduces the *transfer function*, that quantifies the displacements in response to both external forces and thermal noise, in a frequency decomposed manner. Computing transfer functions at low frequencies involves the inversion of an ill-conditioned (nearly singular) matrix, arising from the degrees of freedom related to rigid body rotations and translations. We address this issue by reformulating the inversion of a matrix as a constrained least-squares problem (CLS), where the constraints are obtained from Eckart’s conditions [13, 14]. The solution of the CLS problem yields the transfer functions needed. Section 2.3 introduces the *power spectral density* (PSD) that characterizes both deterministic and stochastic signals in a frequency decomposed manner. Section 2.4 presents a general scheme that utilizes the transfer functions in order to compute the PSDs of the displacements (outputs) from the PSDs of the applied forces and noise excitations (inputs). We describe in Section 2.5 how the mean square fluctuations of node (atom or coarse-grained bead) positions can be computed using transfer functions and PSDs when the network is excited only by noise. This is accomplished by simply integrating the PSDs over all frequencies. We show that the results obtained as such match the ones that are computed using a well known technique [15] that uses the pseudo-inverse of the Hessian of the potential energy function. The equivalence of the two techniques not only validates the constrained transfer function and the PSD approach, but also leads to the characterization of the fluctuations in a frequency decomposed manner. In Section 2.6, we define frequency decomposed SNR as the ratio of the displacement PSDs due to the deterministic external forces (modeling ligand binding) to the ones due to noise alone. Finally, channel capacity is defined and computed based on frequency decomposed SNRs.

### 2.1 Dynamics: Langevin Formulation

We consider *N* particles interacting according to a potential energy profile, *U*, a function of particle positions, 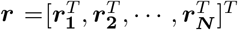, where ***r***_***i***_ = [*x*_*i*_, *y*_*i*_, *z*_*i*_]^*T*^ is the position of the i^*th*^ particle. Each particle may correspond to either an atom or a coarse-grained bead (i.e., a collection of atoms), depending on the resolution of modeling. The potential energy is expanded around the equilibrium (minimum energy) position 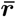 as

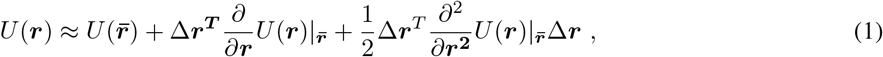

where 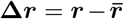 is the displacement from the equilibrium position. The first-order derivative of the potential energy vanishes at the energy minimum, and the constant energy term can be set to zero without any impact on the dynamics. Hence, the potential energy around the equilibrium point is approximated with a quadratic function as

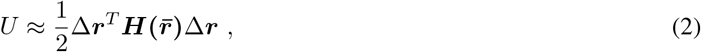

where ***H*** is the Hessian (also known as the force constant matrix), and structured as

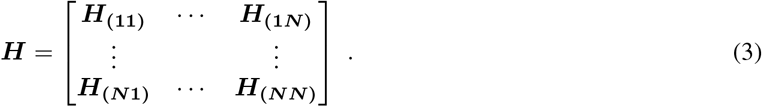

The Langevin formulation describes the protein dynamics in solution [16], as governed by the following coupled set of 3*N* equations written in matrix form

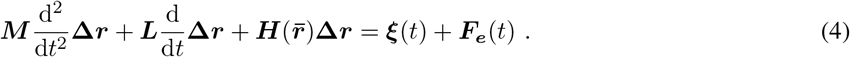

***M*** = diag(***M***_(1)_, ***M***_(2)_, …, ***M***_(*N*)_) is a block diagonal mass matrix. The effect of frictional forces exerted by the viscous fluid is incorporated into the dynamics via ***L***, which is taken to be proportional to the velocity of the particles. ***ξ***(*t*) captures the background noise forces due to random collisions of the Brownian particles. 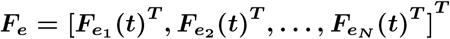 is the external force vector of size 3*N ×* 1. Hydrodynamic shielding effects are ignored, and thus ***L*** = diag(***L***_(1)_, ***L***_(2)_, …, ***L***_(*N*)_) is a diagonal matrix of the friction coefficients. We consider the special case of spherical particles, and the frictional force is assumed to act isotropically on the particles, i.e., ***L***_(*i*)_ = diag(*γ*_*i*_, *γ*_*i*_, *γ*_*i*_). The friction coefficient *γ*_*i*_ for particle *i* with radius *a*_*i*_ is given by Stoke’s law as

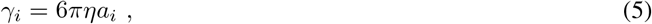

where *η* is the dynamic viscosity, and the hydrodynamic radius is calculated from the volume of the particle, *V*_*i*_, as 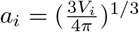.

In the Langevin formulation, the noise and the viscous friction terms are linked to each other via the fluctuation-dissipation theorem [17, 18]. Particle positions continually fluctuate due to the interplay between the noise and the friction in the system, even in the absence of external force excitations, i.e., when ***F***_***e***_ = **0. *ξ***(*t*) in the Langevin equation is a wide sense stationary (WSS) stochastic process. The first and second moments of the noise force exerted on particle *i*, a component of ***ξ***(*t*), is given by [16]

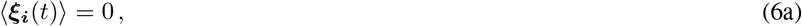

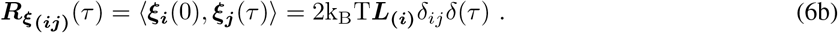

Above, ⟨·⟩ denotes an average with respect to the distribution of the realizations of ***ξ*(*t*)**. Due to ergodicity, the time and ensemble averages are equal. ***R***_***ξ***_(*τ*) = E[***ξ***(0)***ξ***(*τ*)^*T*^] is the auto-correlation matrix of the random force ***ξ***(*t*), of size 3*N* × 3*N*. The Dirac delta function *δ*(*τ*) indicates that there is no correlation between different time samples of the random forces. The Kronecker delta function *δ*_*ij*_ indicates that random forces acting on different particles are uncorrelated. Therefore, ***R***_***ξ***_(*τ*) is a diagonal matrix. *i* and *j* represent particle indices, *i, j* = 1, 2, …, *N*.

### 2.2 Transfer Function

The input-output characteristics for a linear and time-invariant (LTI) system can be described through transfer functions. In the system of interest, ***T*** represents a matrix of transfer functions from the inputs, the external forces or the noise acting on the system represented by ***F***, to the output, displacement deviations **Δ*r***

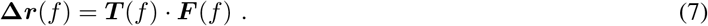

where *f* is the frequency of the input and also the output. LTI systems excited at a single frequency *f* produce a response that also has a single frequency component at the same frequency. Furthermore, LTI systems satisfy the superposition property: The sum of the responses to two different inputs is equal to the response to the sum of the inputs. With ***F*** set to either the noise input ***ξ***(*t*) or the force excitation ***F***_***e***_(*t*), Equation 4, the equation of motion, is written in the frequency domain as

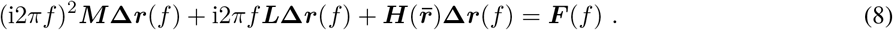

where 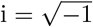. Thus, complex-valued transfer function ***T*** (*f*) is defined as

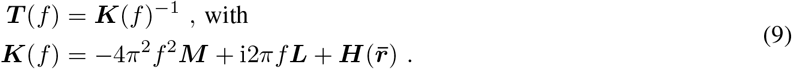

Transfer functions in the frequency domain were also utilized in our recent previous work on probing the perturbation-response dynamics of proteins [10].

#### 2.2.1 Constrained Transfer Function

The Hessian ***H*** is a singular matrix with a rank deficiency of six, due to the degrees of freedom arising from rigid-body rotations and translations of the molecule. Therefore, the transfer function computation is ill-conditioned at low frequencies, and not possible at zero frequency. To eliminate the six degrees of freedom, the following set of translational and rotational Eckart’s conditions are introduced [13, 14]

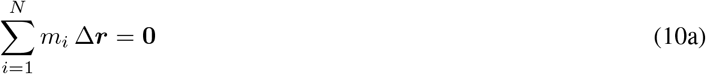

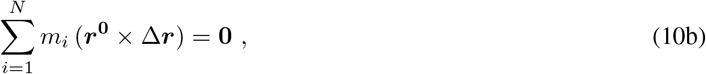

where × indicates a vector product, *r*^0^ = *r*_*i*_ − *r*_*com*_ is the displacement from the center of mass, *r*_*com*_, and *m*_*i*_ is the mass of the i^*th*^ particle. Equation 10 can be recast in matrix form as follows

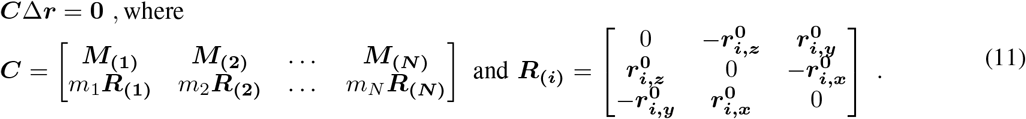

With the incorporation of these constraints, the inversion of a nearly rank-deficient matrix is replaced with the solution of the following constrained least-squares problem

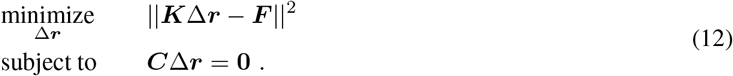

A Lagrange multiplier-based solution of the above problem yields the following (Please see Section S.2 for the derivation.)

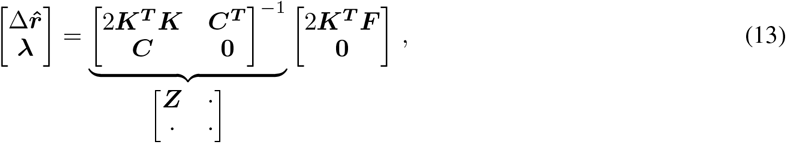

where ***λ*** =[*λ*_1_,*λ*_2_,*…, λ*_6_]^*T*^ are the Lagrange multipliers. The transfer function matrix incorporating the Eckart constraints, denoted by ***T***_***c***_, can be readily constructed using the first 3*N* rows and columns of the matrix that is obtained as the inverse of an augmented matrix shown in the RHS of Equation 13, denoted by ***Z***. Thus,

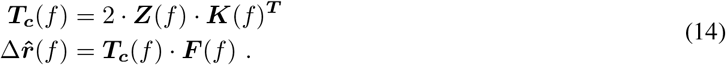

In the rest of the paper, whenever appropriate, the singularity or ill-conditioning problem in matrix inversion is addressed as above.

### 2.3 Power Spectral Density (PSD)

The power spectral density describes the content of a signal in a frequency decomposed manner. It can be computed either directly from the squared amplitudes of the frequency domain representation (for deterministic signals), as in Equation 15a below, or as the Fourier transform of the auto-correlation function (for stochastic signals), as in Equation 15b), due to the Wiener-Khinchin theorem [19]. ***S***_***X***_ is the PSD of a signal *X*

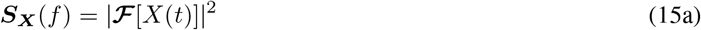

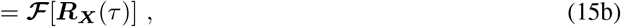

where **ℱ** denotes the Fourier transform, defined with 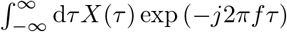 for the signal *X*. Equation 15a is used for external force excitations that are deterministic, Equation 15b is used for noise sources and stochastic fluctuations.

#### 2.3.1 PSD of Noise Sources

The PSD of the noise force acting on the *i*^*th*^ particle can be determined by inserting the autocorrelation 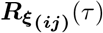 defined in Equation 6b into Equation 15b

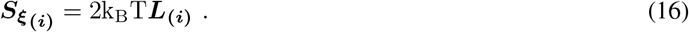

The above PSD is constant over the frequency spectrum, corresponding to *white noise*.

#### 2.3.2 PSD of External Force Excitations

A deterministic external force that acts on certain nodes mimicks the ligand binding event. Although the proposed method does not impose any restriction on the form of the external force ***F***_***e***_, we consider it as a dynamic, sinusoidal perturbation whose frequency is swept over a certain range, allowing us to quantify the effect of perturbation frequency. The external force applied to particle *j* is in the form

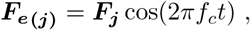

where ***F***_***j***_ is a 3 ×1 vector that contains the force components along the coordinate axes applied on the j^*th*^ particle, and *f*_*c*_ is the frequency.

The PSD of this sinusoidal force at the input site is given by

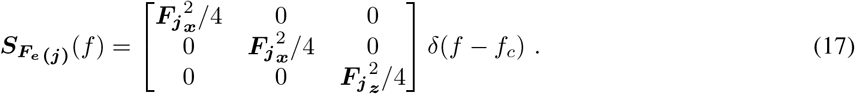

### 2.4 From Excitation PSDs to Displacement PSDs

For multi-input multi-output LTI systems, the cross spectral density matrix of the outputs, ***S***_***Y***_, can be calculated as follows by using the cross spectral density matrix of the inputs, ***S***_***X***_, and the transfer function ***T*** (please see [20] for the derivation)

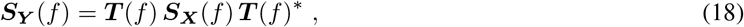

where .^*^ denotes complex conjugate. When the input is the force excitation (either due to noise or external force), and the output is the fluctuations/displacements of node positions, the above relationship can be written using the constrained transfer function defined in Equation 14 as

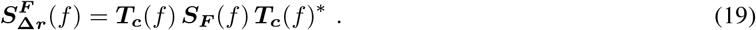

We separately compute the PSDs of the output due to noise and deterministic excitations by first setting ***F*** = ***ξ*** and then ***F*** = ***F***_***e***_. We note that 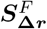 is a full matrix. The diagonal entries represent the spectral densities, whereas the off-diagonal entries are the cross-spectral densities. We focus on the diagonal entries in this study. Hence, we introduce the following notation

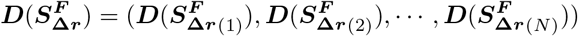

for the 3*N* × 1 vector formed from the diagonal elements of 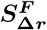.

### 2.5 Characterization of Equilibrium Fluctuations based on the PSDs of Displacements due to Noise

The partition function of the system, whose potential energy profile is given in Equation 2, can be calculated over all possible displacements with 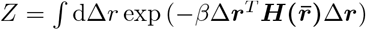, where *β* = 1*/*k_B_*T*, k_B_ is the Boltzmann’s constant, and T is the absolute temperature. The correlations of the fluctuations between sites *i* and *j* can be obtained from this partition function as [21]

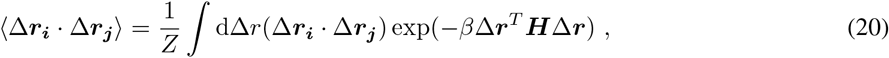

which can be simplified to

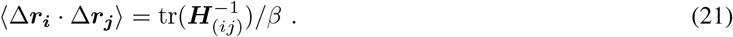

Thus, mean square fluctuations for particle *i* is

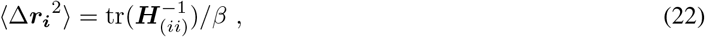

where ***H***_(*ij*)_ corresponds to the 3 × 3 submatrix for particles *i* and *j*, and tr denotes the trace. Since the Hessian is singular as discussed before, its pseudo-inverse (***H***^**+**^) is computed through the use of the non-zero eigenvalues, ***µ***_*k*_, and the corresponding eigenvectors, ***u***_*k*_, as follows

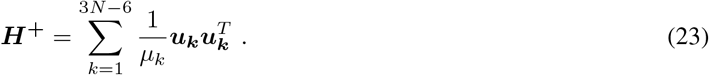

The mean square fluctuations computed as such are proportional to the B factors determined from X-ray crystallography, also known as temperature factors [15], denoted by *B*:

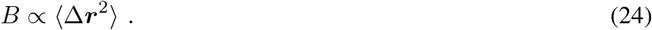

The total power in a WSS stochastic signal *X* can be expressed in terms of the variance of the signal, i.e., zero time-lagged auto-correlation, which is also given by the integral of its PSD over all frequencies, as below

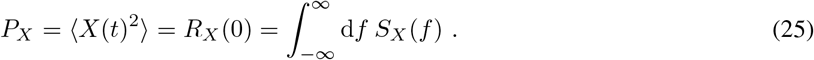

In the absence of external force excitations, the mean square fluctuation of node *i* can be thus computed as the sum of integrals of the displacement PSDs along the coordinate axes as

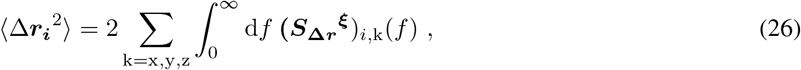

where the factor 2 is due to the symmetry of the two-sided PSD for negative and positive frequencies. This PSD-based technique thereby offers a previously overlooked route to computing mean square fluctuations. Previously published studies mostly focused on the frequency distribution of the B factors at a set of discrete frequency points, known as the normal mode frequencies. The PSD-based approach allows the frequency decomposed analysis of B factors over any continuous frequency range of interest.

### 2.6 Signal-to-Noise Ratio (SNR) and Channel Capacity

We define SNR as the ratio of the PSD of the displacements measured at the output due to external force excitations alone and the displacements due to only noise:

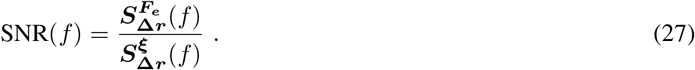

According to the integral-form of the Shannon-Hartley theorem [12], the channel capacity that can be attained over a noisy communication channel is defined as

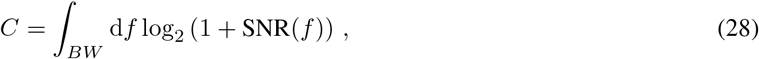

where the integral is computed over a frequency band *BW* of interest. We choose [0, ∞) as the frequency interval. Then, the channel capacity between the input node *j* and output node *i* can be calculated as the sum of the channel capacities along the coordinates axes

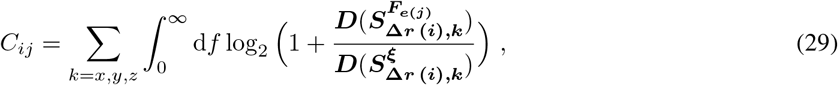

where *F*_*ej*_ indicates that the deterministic force excitation is applied to node *j* only but results in dynamic displacements at all the others. In computing 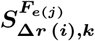, the strength of the input excitation is kept constant while its frequency is swept. The strength of the resulting response will vary not only as a function of the output node but also over the frequency range, as determined by the internal network dynamics. Thus, while information (signal) transmission between two particular nodes may be inefficient at a certain frequency, it may be enhanced at another. On the other hand, random noise excites all nodes simultaneously, at all frequencies with equal strength, and it is shaped by the network dynamics resulting in fluctuations at all nodes with a colored frequency composition as opposed to the white noise excitation. The extent of fluctuations due to noise is also a function of the output node. We emphasize here the distinction between signal transmission and noise propagation through the network. While the signal permeates throughout the network from a single entry point, noise enters from everywhere. Thus, the frequency decomposed profiles of node displacements due to either the signal or the noise alone may be quite different, resulting in a colored frequency composition for SNR with possibly non-monotonic behavior.

## 3 Methods

Figure 1 presents a pictorial overview of the methods described in Section 2. The protein is represented as a network, and the protein-solution interactions are modeled as random noise forces that act on every node. In contrast, the effect of the ligand is captured by a deterministic, oscillatory external force, which is applied to a single node (or simultaneously to a set of nodes), denoted as the input node(s). The goal is to examine how the effect of the external force excitation is transmitted to the output node in the presence of all of the noise sources. The random noise is white, i.e., has a constant PSD over the frequency interval of interest. The magnitude of the external force is held constant as a function of frequency. The PSDs measured at the output node are calculated using the input PSDs and the frequency-dependent transfer functions, quantifying the displacements in response to both deterministic and noise excitations. We underline that although noise forces are applied to all of the nodes, noise PSD only for the input node, upon which the external force is applied, is shown in the pictorial overview for brevity. The methods proposed in this paper enable a frequency decomposed analysis, by sweeping the perturbation frequency in a frequency range of interest. The gray bars in the PSDs represent frequency sweeping, while the highlighted bars correspond to the perturbation frequency that is currently under investigation. The PSD plots at the output show the frequency-dependent characteristics. SNR is calculated using the output PSDs, by simply taking the ratio of the PSDs due to deterministic and noise excitations at each frequency. The channel capacity is computed via an integral of the SNR over a frequency band. In Figure 1, the color green represents noise and red is used for the external force. Yellow and purple are used to distinguish the input and output nodes, respectively.

**Figure 1:**
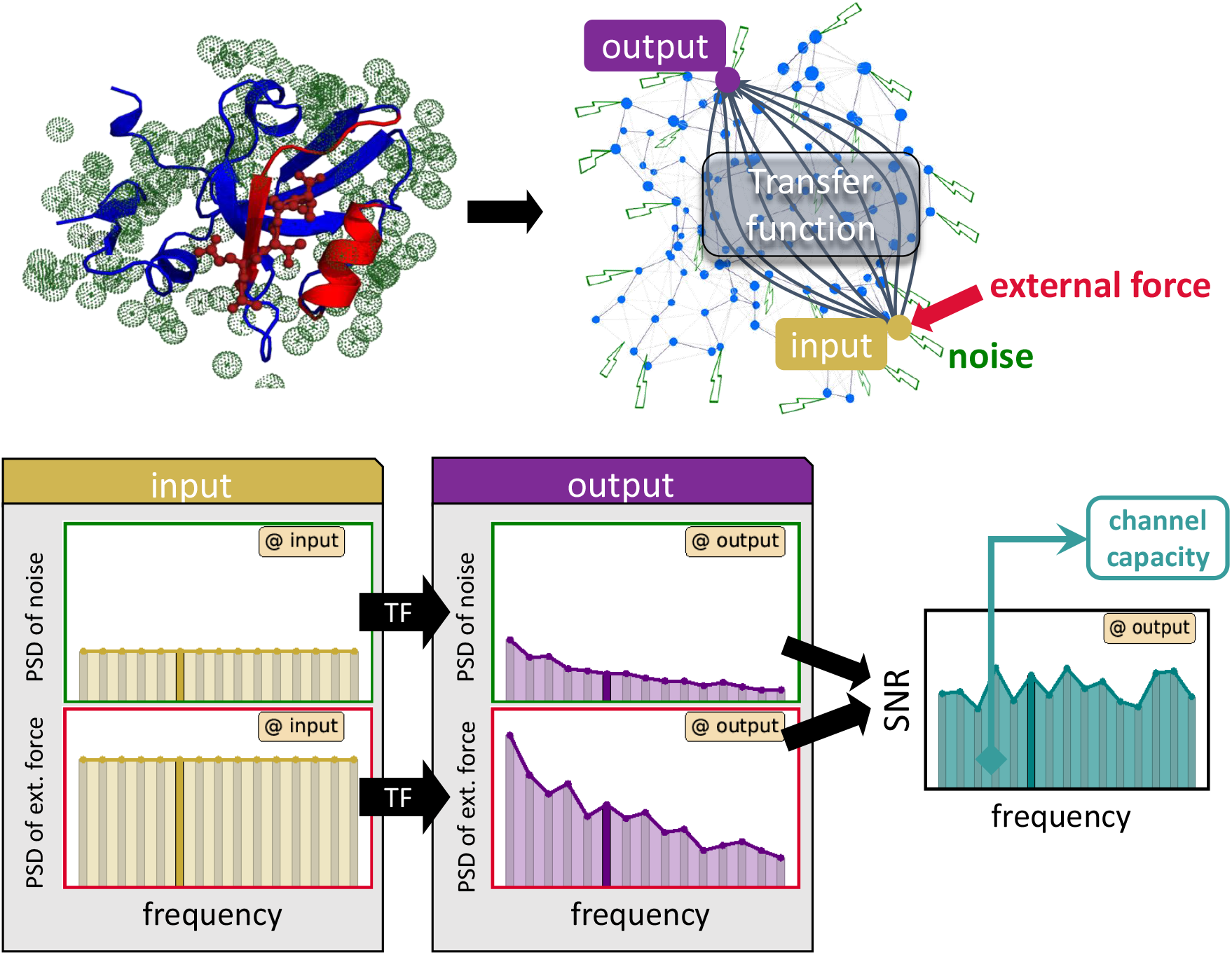
A simplified pictorial overview of the work flow. The signal permeates throughout the network from a single entry point, the input, and converges at the output. Noise enters the network from everywhere and every noise component has an impact at the output. Although the noise forces act on every node, only some of them are shown in the figure to reduce clutter. The network was drawn using the NAPS web-server provided in [22].

An algorithmic summary of the proposed methods is presented in Figure 2. The top flow line schematically shows the steps for obtaining the B factors from the Hessian. We use two alternate methods for Hessian construction, described in Section 3.1. One approach relies on molecular dynamics (MD) while the other is based on elastic network models (ENMs). In Figure 2, the two parallel flow lines below are for the operations involved in obtaining the output displacement PSDs from the input PSDs, including the computation of the constrained transfer functions, both for deterministic excitations and noise. As an alternative to direct inversion of the Hessian, the PSDs of the displacements due to noise not only lead to the B factors, but also enable the frequency decomposed analysis of the equilibrium fluctuations. SNRs are computed as the ratio of the displacement PSDs due to external force excitations and to those due to noise, subsequently integrated to obtain the channel capacities.

**Figure 2:**
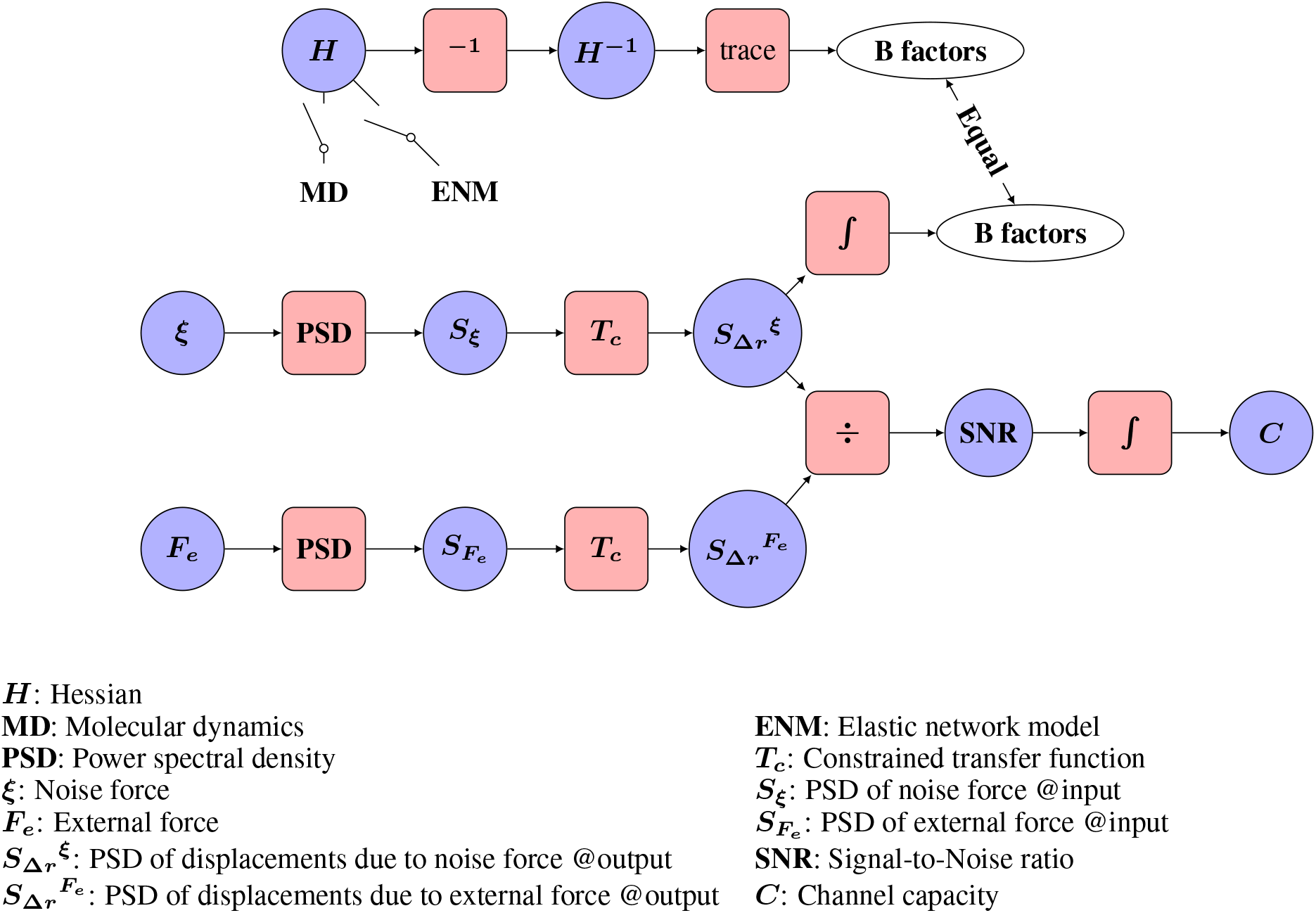
Schematic of the work flow.

### 3.1 Construction of Hessian

The second order partial derivatives of the potential energy function (evaluated at the equilibrium point), captured in the Hessian ***H***, that are used in the Langevin formulation described in Section 2.1, are obtained following two alternate approaches: (i) from molecular dynamics simulations (MD) and (ii) based on elastic network models (ENMs).

#### 3.1.1 Hessian from MD

In the first step, positional fluctuation trajectories around a reference or equilibrium structure are calculated. Next, the covariance matrix is formed based on the time averages over the trajectories as

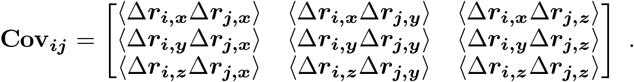

**Cov**_***ij***_ above is a 3 ×3 matrix of the covariances between the *x, y, z* components of the positional deviation vectors **Δ*r***_***i***_ and **Δ*r***_***j***_ for nodes *i* and *j*. The overall covariance matrix **Cov** is then formed as a block matrix of size 3N× 3*N*, where the *ij*^*th*^ block is set to **Cov**_***ij***_. The Hessian matrix is in fact equal to the scaled inverse of the covariance matrix, ***H***^MD^ = k_B_T **Cov**^−1^ [23], where k_B_ is the Boltzmann’s constant and T is the absolute temperature.

#### 3.1.2 Hessian from ENMs

In ENMs, the protein is represented as a mass-spring network, resulting in a simplified representation of the potential energy function. The nodes of the network, typically chosen as the *C*_*α*_ atoms of the residues, are interconnected by springs. The nominal node positions are determined based on a reference structure, which is usually experimentally determined. The interactions are considered for the residue pairs within a pre-defined cutoff distance of each other. The reference structure is assumed to correspond to the global minimum of the potential energy function by construction, hence no energy minimization is needed. Gaussian Network Models (GNM) [24] and Anisotropic Network Models (ANM) [25] are two subclasses of ENMs. GNMs take into account only the magnitude of the fluctuations, while ANMs capture the directionality as well. The potential energy function for ANMs can be written as

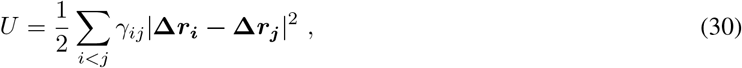

where *γ*_*ij*_ is the spring constant between residues *i* and *j*, and| ·| denotes the *L*_2_ norm (Euclidean length). The Hessian of the potential energy is constructed as

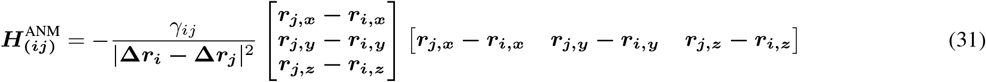

for *i* ≠ *j* where (·)_*x,y,z*_ denotes the Cartesian coordinates along the corresponding coordinate axes, and the diagonal block for *i* = *j* is set to

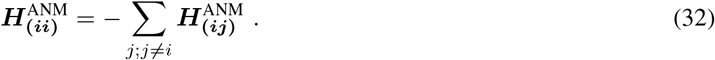

ENMs have been successful in predicting the residue fluctuation profiles of globular proteins [26].

### 3.2 Molecular Dynamics Simulation Details

We considered two distinct systems for simulations, the PDZ3 domain with (PDBID:1be9) and without (PDBID:1bfe) the associated ligand. Starting from the PDB structures, missing atoms were added to the structure using the PyMOL mutagenesis tool [27]. Each structure was then solvated with sufficient number of water molecules to “fill” a cubic box with sides of length 6.6 nm (8804 and 8845 water molecules for the system with and without the ligand, respectively), using the *gmx solvate* command in the GROMACS package [28]. Additionally, the minimum number of ions were added to neutralize the system (2 Na ions for the system with the ligand and 1 Na ion for the system without the ligand). The Amber99SB-ILDN force field [29, 30] was used to model the protein interactions, while the TIP3P model [31] was used to model the water interactions.

All simulations were performed with the Gromacs 5.0.8 simulation suite [28] with a 2 fs integration time step, while using the LINCS [32] algorithm to rigidly constrain all bonds that involve H atoms. All simulations employed the Gromacs leap-frog integrator [33] and periodic boundary conditions [34]. Electrostatic interactions were treated with the particle mesh Ewald method [35], using the default Gromacs parameters. Short-ranged van der Waals interactions and also the real space contribution to the electrostatic interactions were truncated at 1.2 nm in all simulations. After energy minimization, each system was annealed from 0 to 300 K over a 500 ps time frame, followed by a 5 ns equilibration simulation in the *NV T* ensemble using the Berendsen thermostat [36] with a temperature coupling constant of 0.5. Subsequently, an *NPT* equilibration was performed using the Berendsen thermostat and barostat with coupling coefficients of 0.1 for both and a compressibility of 4.5 × 10^−5^ bar^−1^ (corresponding to the compressibility of water at 1 atm and 300 K). The average volume from this simulation (corresponding to box side lengths of 6.56797 nm and 6.56372 nm for the systems with and without the ligand, respectively) was used to select an initial structure for the production simulation. Production simulations were performed for 100 ns in the *NPT* ensemble, using the velocity rescaling thermostat [37] with a temperature coupling coefficient of 0.1 and the Parrinello-Rahmen barostat with a pressure coupling coefficient of 2.0. Configurations, velocities, and forces were saved every 2 ps for subsequent analysis. We characterized the local impact of the ligand by calculating the average force that all atoms of the ligand exert on each of the C_*α*_ atoms of the binding pocket residues (residues [320 − 328] and [371 − 380]).

### 3.3 External Force Excitations

Figure 3 presents a schematic view of the two different techniques we used in order to probe allosteric behavior, namely, *per residue scan* and *binding pocket (BP) excitation*. An external force is applied to only one residue in the first case, whereas multiple residues are perturbed simultaneously in the latter. With *per residue scan*, the goal is to identify the residue pairs that are likely to interact allosterically. Each residue is separately perturbed with a sinusoidal force whose frequency is swept from 0 to the maximum of the normal mode frequencies. The force in this case has a unit magnitude but numerous directions are sampled on a spherical grid. The channel capacities from the input to the rest of the residues are computed for each force direction, and a distinct direction for every output residue that maximizes its capacity is identified. *BP excitation* models ligand binding to the protein in a more realistic manner by simultaneously exciting multiple residues that are previously known to be in a certain binding pocket. The responses of all of the remaining residues are monitored. Instead of sampling the force directions on a grid, the relative amplitudes and the directions of the external forces that act on the binding pocket residues are directly obtained from MD simulations as described in detail in Section 3.2, while the sinusoidal form and the frequency range are as in *per residue scan*. The channel capacities calculated as such are normalized (in a min-max sense such that the range of the channel capacity values from the minimum to the maximum are linearly transformed into the range 0 to 1) in order to attain a distribution of capacities within a protein. In the normalization process, the values above a predefined threshold are set to 1, while the remaining values are scaled to be in the interval [0, 1].

**Figure 3:**
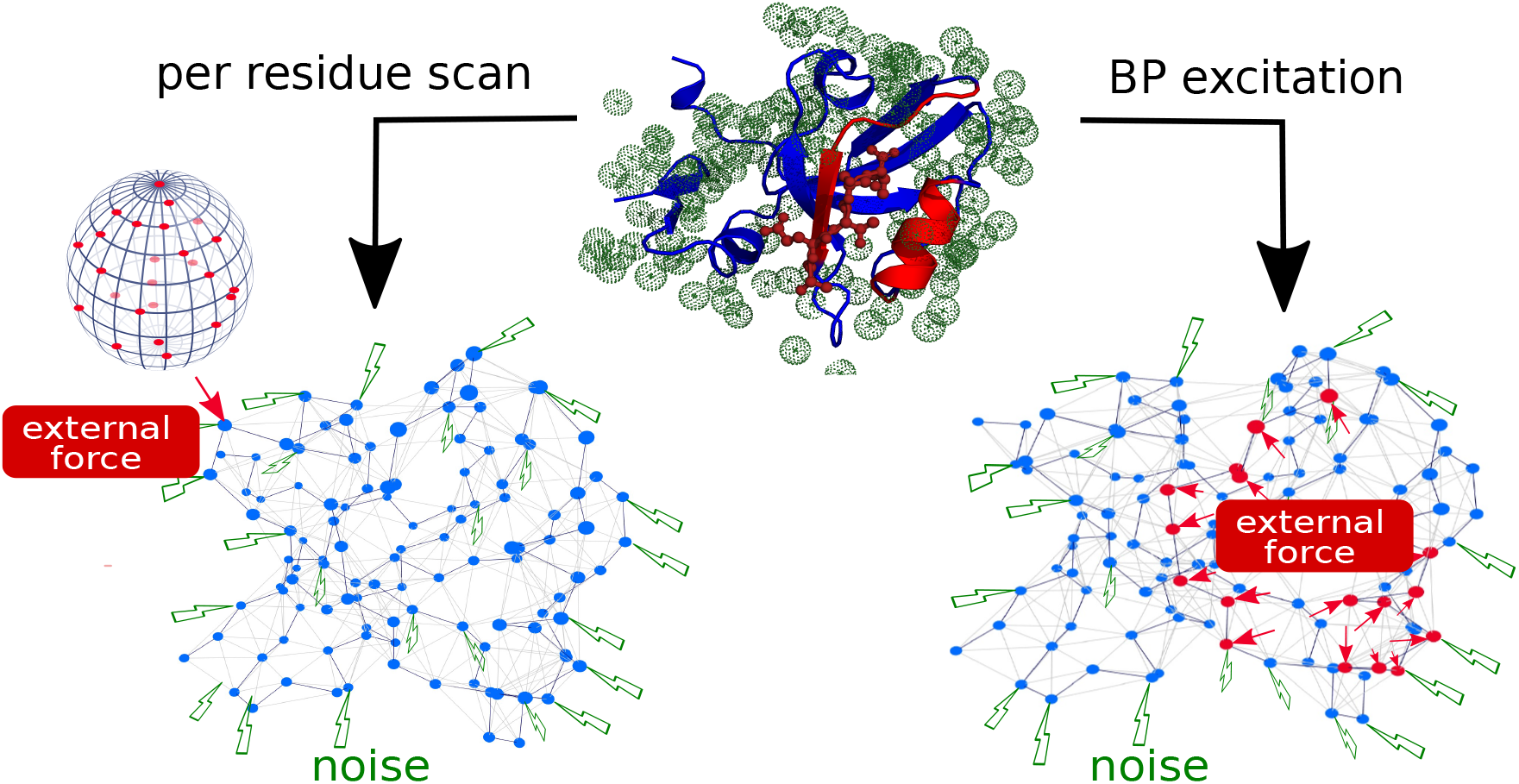
Schematic view of the two different techniques used in order to probe allosteric behavior.

### 3.4 Parameters

Residues are represented by their C_*α*_ atoms. A 200-point Gauss-Legendre quadrature scheme from NumPy package is used to numerically calculate the integrals [38, 39]. *η*, which is utilized in the friction coefficient calculation in Equation 5 and subsequently in Equation 4, is taken as the dynamic viscosity of water at 310 K, that is equal to 6.7807 · 10^−4^ Pa · s. Masses and volumes of the residues are retrieved from [40] and [41], respectively, also listed in Table S1. However, as shown in Section S.3, the mean square fluctuations are independent of the friction coefficient values due to the link between noise PSDs and friction in the formulation in Equation 4. We utilized ANMs with a cutoff distance of 15 Å and the same spring constant for all interactions. In *per residue scan*, the force directions are sampled from a 25 point equally-spaced spherical grid. The values above 90% of the maximum channel capacity value are normalized to a value of 1. In the *BP excitation*, the force exerted by the ligand on the binding pocket residues are determined from MD simulations of the holo form as described in Section 3.2. The rotation matrix calculated for the superposition of the apo and holo forms is then applied to this external force vector so that it can be used as a force excitation for the apo form.

## 4 Results

We demonstrate the utility of the proposed methods on a well-studied PDZ3 protein (the third PDZ domain of PSD-95) that is known to display dynamic allostery, i.e., allostery without major structural changes. PDZ domains are one of the most abundant and evolutionarily conserved protein-protein interaction modules that take part in numerous cellular and biological functions, including dimerization and recognition of specific sequences of C terminus tails of other proteins [42]. The conserved structures in PDZ domains are six beta strands: *β*1− *β*6, and two alpha helices: *α*1 and *α*2. Some of the PDZ domain proteins have additional structures, which have been proposed to affect ligand binding. PDZ3 has *α*3, *β*7, and *β*8 extensions [43]. It was shown that the removal of the *α*3 domain results in a 21 times decrease in the binding affinity of PDZ3 [44], without a significant conformational change. This points to the allosteric nature of the PDZ3 domains, as well as to the entropic (as opposed to structural) nature of this allosteric behavior. The aligned structures of the 110-residue-long apo and holo forms of PDZ3 (PDB ID’s 1bfe and 1be9, respectively) are shown in Figure 4. The ligand, which is represented in ball-and-stick form, is a 5-residue-long C-terminal segment of the Cysteine-rich PDZ-binding protein CRIPT. The binding pocket lies between the *α*2 − *β*2 regions, and is comprised of residues [320− 328] and [371 −380]. Figure S1 presents the sequence and secondary structure assignment information.

**Figure 4:**
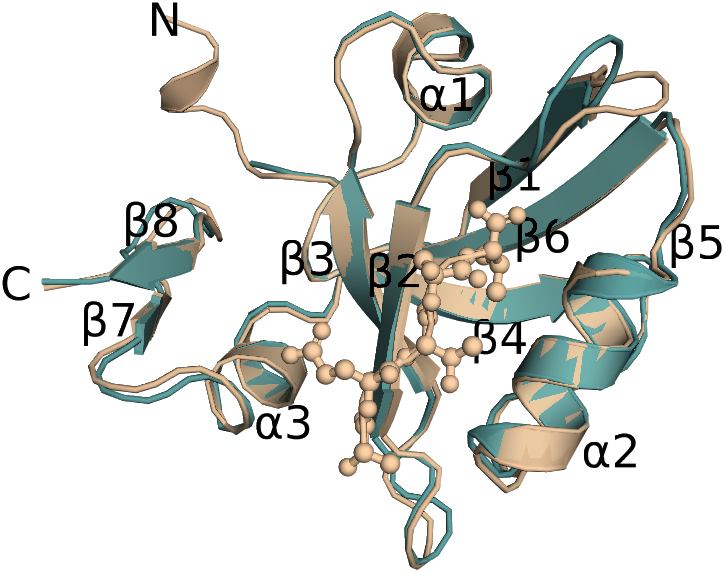
The aligned structures of the apo and holo forms of the PDZ3 protein. Wheat (teal) color is for the holo (apo) form. The ligand is represented in ball-and-stick format. PyMOL was used for the visualization [27].

First, the analysis techniques proposed in this paper are validated in Section 4.1 by examining the displacements due to noise forces alone. This verification is based on the equivalence of the B factor values obtained via the proposed work flow and previously available computational techniques. Since the proposed techniques emphasize frequency domain representations, the frequency distribution of equilibrium fluctuations is obtained as an intermediate step. Then, in Section 4.2, we present frequency-decomposed SNR and channel capacity based analyses in the presence of external perturbations using both the *per residue scan* and *BP excitation* scenarios.

### 4.1 Without External Force: Equilibrium Fluctuations

Mean square fluctuations (*MSFs*), which are proportional to experimentally measurable B factors, can be directly calculated from the pseudo-inverse of the Hessian of the potential energy function (please refer to Equations 22 and 24). We proposed an alternative frequency domain method to compute these equilibrium fluctuations by integrating the power spectral densities over the whole frequency range. The proposed scheme is based on the (linearized) Langevin formalism (please see Equation 4), which uses ***H***. Hence, the *MSFs* obtained from the pseudo-inverse of ***H*** and the *MSFs* from the proposed method should correspond to each other. Here, we consider ***H*** obtained both from MD (***H***^MD^) and also from ANM (***H***^ANM^) (see Section 3.1 for details). Figure 5 presents the B factors obtained for the apo form (PDBID:1bfe): from (i) the direct pseudo-inverse approach, (ii) the proposed PSD-based scheme, and (iii) experimentally measured values. The orange and green lines present B factors obtained from ***H***^MD^ and ***H***^ANM^, respectively. The blue and red lines represent B factors calculated from the integral of the PSDs both with ***H***^MD^ and ***H***^ANM^, respectively (please refer Section 2.5 for the details of the method). The perfect match between the proposed frequency domain scheme and the pseudo-inverse-***H*** technique confirms the validity of the proposed work flow that utilizes the transfer functions with constraints introduced to overcome the singularity problem, and PSD integrations. Moreover, the agreement of the B factors between ANM and MD (except for the residues [379 −381]) based Hessians justifies the use of simplified models for fluctuation analyses around an equilibrium structure. The black line in Figure 5 corresponds to the experimental B factors, which are reported in the associated PDB file. The secondary structure information is displayed at the bottom. The frequency decomposition of the *MSFs* are shown in Figure 6 for ***H***^MD^ and ***H***^ANM^. (Please see Figure S4 for the side views). In both cases, the low frequencies contribute significantly more to the total *MSFs*. Also, there is an apparent trend: PSDs decrease as frequency increases, yet with varying rates for different residues. The PSDs obtained from ***H***^MD^ display a more rapid decrease starting from low frequencies, whereas for ***H***^ANM^, this decrease starts at higher frequency values. Although the difference in total *MSFs* for the two Hessians is relatively small, as seen in Figure 5, the spectral decompositions exhibit substantial differences.

**Figure 5:**
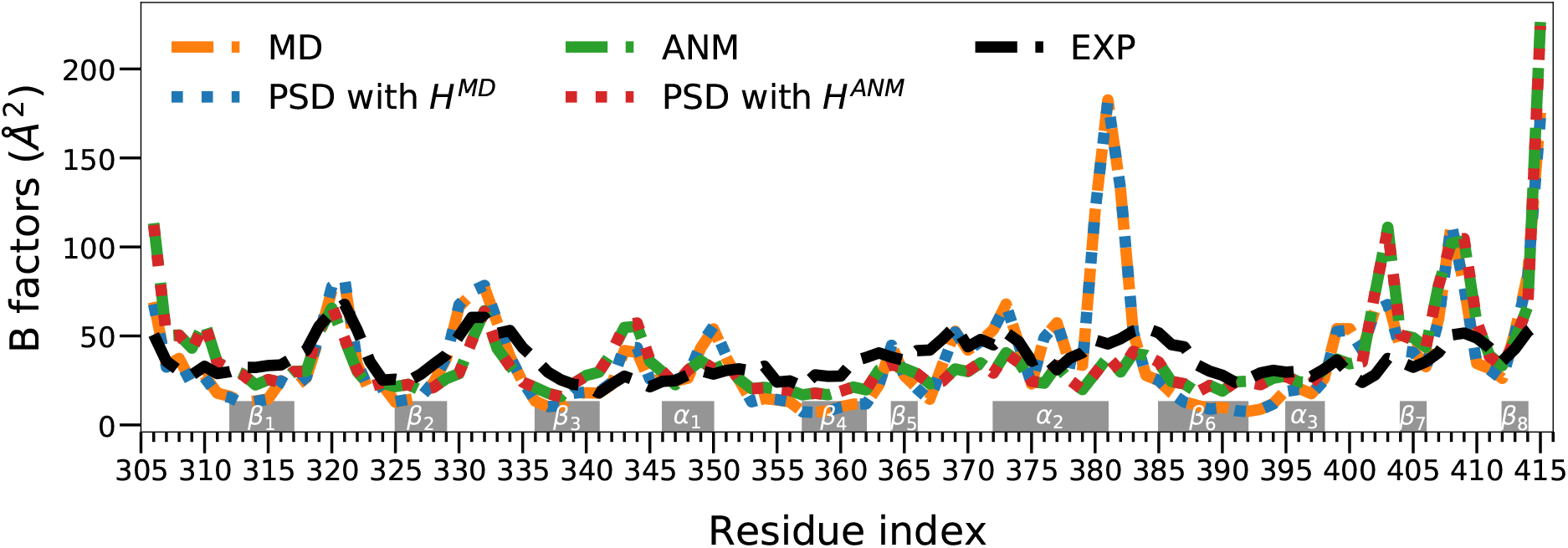
B factor values that are calculated with various methods for the apo form (PDBID:1bfe). The blue (red) lines show the values calculated via power spectral density integrations using ***H***^MD^ (***H***^ANM^). The orange (green) line is calculated directly from the pseudo-inverse of ***H***^MD^ (***H***^ANM^). The black line shows the experimental values. The values are scaled to correspond to the experimental ones.

**Figure 6:**
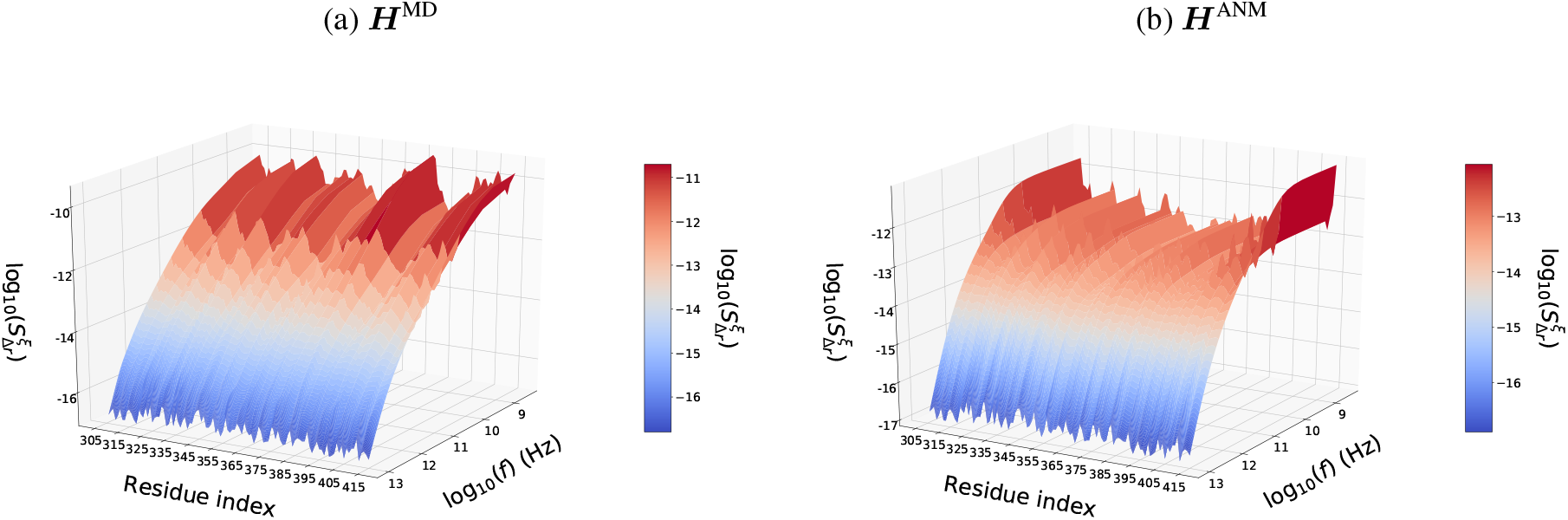
PSD profile of the equilibrium fluctuation with (a) ***H***^MD^ and (b) ***H***^ANM^.

### 4.2 External Force Excitations

#### 4.2.1 Per residue scan

Figure 7 presents the capacity values obtained with *per residue scan* for ***H***^MD^. The normalized channel capacity values presented in panel (a) suggests a coupling between the first several residues closest to the C terminal and (*β*_7_ & *β*_8_) domains, as well as between *β*_1_ and *β*_6_, *β*_2_ and (*β*_3_ & *β*_4_), *β*_5_ and *β*_6_, and *β*_7_ and *β*_8_ domain pairs. Due to the fact that high channel capacity is expected between residue pairs that are close to each other in space, the capacity values are distance-weighted to detect the long-range interactions more clearly, as shown in Figure 7(b). The pairwise distance values that are used in this weighting procedure are calculated at the equilibrium point (please see Figure S2 for the pairwise distances). Despite being distant, the residue pairs that are found to be linked are listed in Table 1. The motions of SER 320 - SER 408, SER 409, and ASN 415 appear to be coupled. The results with ***H***^ANM^ for *per residue scan* (as well as for *BP excitation*) are presented in Figure S3.

**Figure 7:**
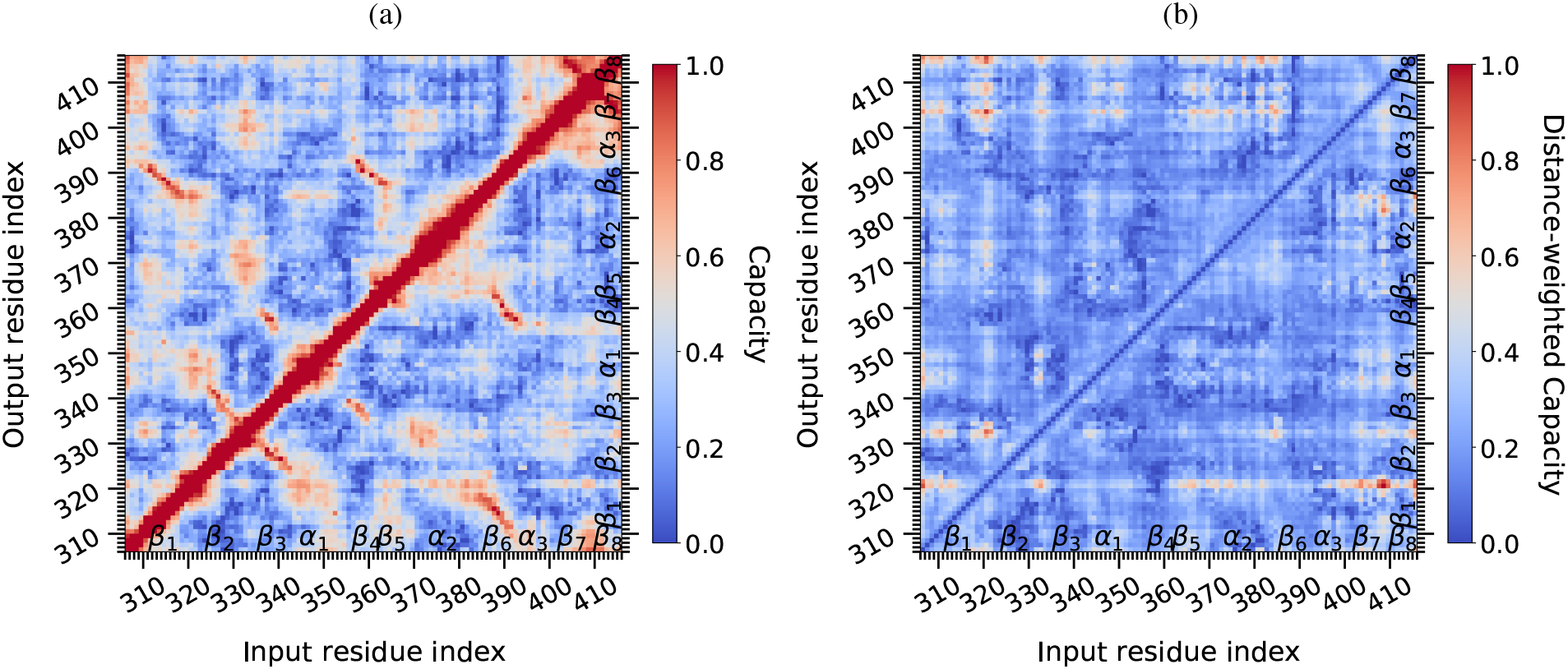
Per residue scan with ***H***^MD^. (a) Normalized, (b) Distance-weighted and normalized channel capacity values.

**Table 1:** Residue pairs that are located far apart in space, yet have high channel capacity values that are identified by *per residue scan*, with ***H***^MD^.

| Input | - | Output |
| --- | --- | --- |
| ASP 306 | - | SER 320, THR 321 |
| SER 320 | - | ASP 332, ASN 403, THR 414, ASN 415 |
| ASN 407 | - | SER 320, THR 321 |
| SER 408 | - | SER 320, THR 321, ASN 381 |
| SER 409 | - | SER 320 |
| ASN 415 | - | SER 320, THR 321 |

#### 4.2.2 Binding pocket (BP) excitation

Figure 8 presents the channel capacity values obtained with *BP excitation* for ***H***^MD^. Panel (a) presents the distance-weighted and normalized channel capacity values, while panel (b) presents a complementary visualization of the 3D protein structure without applying the weighting (The capacity values utilized in panel (b) are shown in Figure S5 in the same format of panel (a)). Please note that the residues that are located in the BP, which are represented by gray boxes in the left panel, are excluded from the normalization procedure since the external forces are directly applied to them. According to the top panel, ASP 306, HIS 317, ALA 343, GLY 344, GLY 383, GLN 384, THR 385, ASN 403, ASN 407, SER 408, SER 409, THR 414, and ASN 415 are identified to have the potential to display the most significant allosteric response to the ligand-binding event.

**Figure 8:**
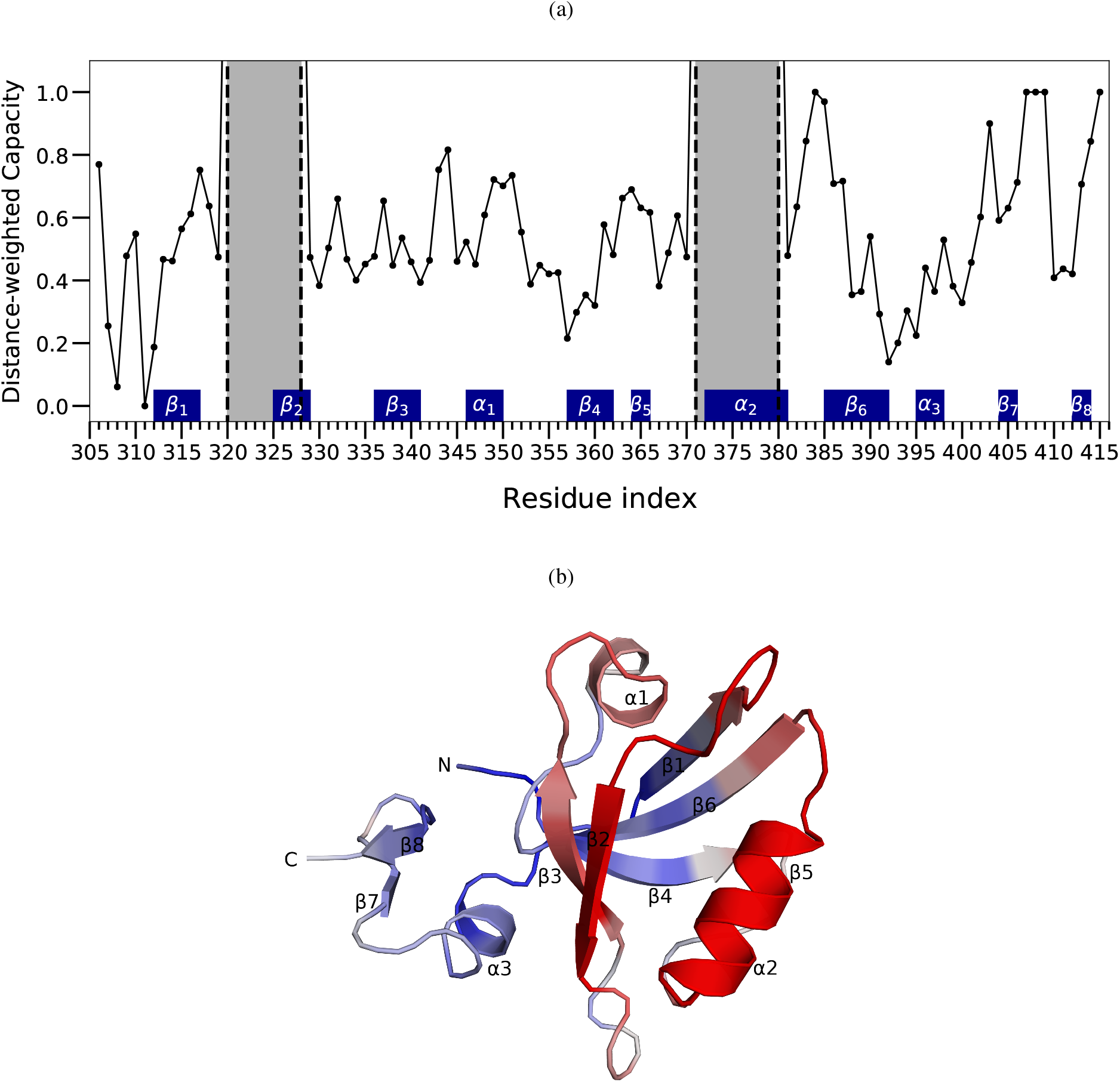
BP excitation with ***H***^MD^ (b) Distance-weighted and normalized channel capacity values. The gray area corresponds to the binding pocket residues upon which the force is applied. (b) Apo protein structure (PDBID:1bfe) is colored according to the capacity values (without distance weighting): red indicates the highest capacities and blue is for the lowest.

Figure 9 presents the residues that have been reported as *critical* using various computational and experimental techniques, which are also listed in Table S2. The normalized channel capacity values (without distance-weighting) obtained from *BP excitation* are displayed as color coded at the bottom row. As in the previous studies, the BP residues are excluded from the analysis. As can be seen from the figure, there is little agreement among the methods that aim to identify the critical residues involved in the PDZ3 protein. Nonetheless, there are several residues that most of the methods agree on, such as GLY 329, ILE 338, ILE 341, ALA 347, LEU 353, VAL 362, and VAL 386. These residues (with the exception of LEU 353) do have a high capacity value based on the results obtained with our channel capacity analysis method. In addition to those listed here, the rest of the residues that have high capacity values were also reported to be critical by at least several previous techniques. Even though these findings point to the success of the proposed method, it is not possible to assess the degree of accuracy with respect to a golden reference due to the lack of consistency among previous methods. Nevertheless, we believe that the proposed channel capacity technique provides a fresh approach for deciphering the mechanisms of allostery, and could help detect hidden allosteric interactions.

**Figure 9:**
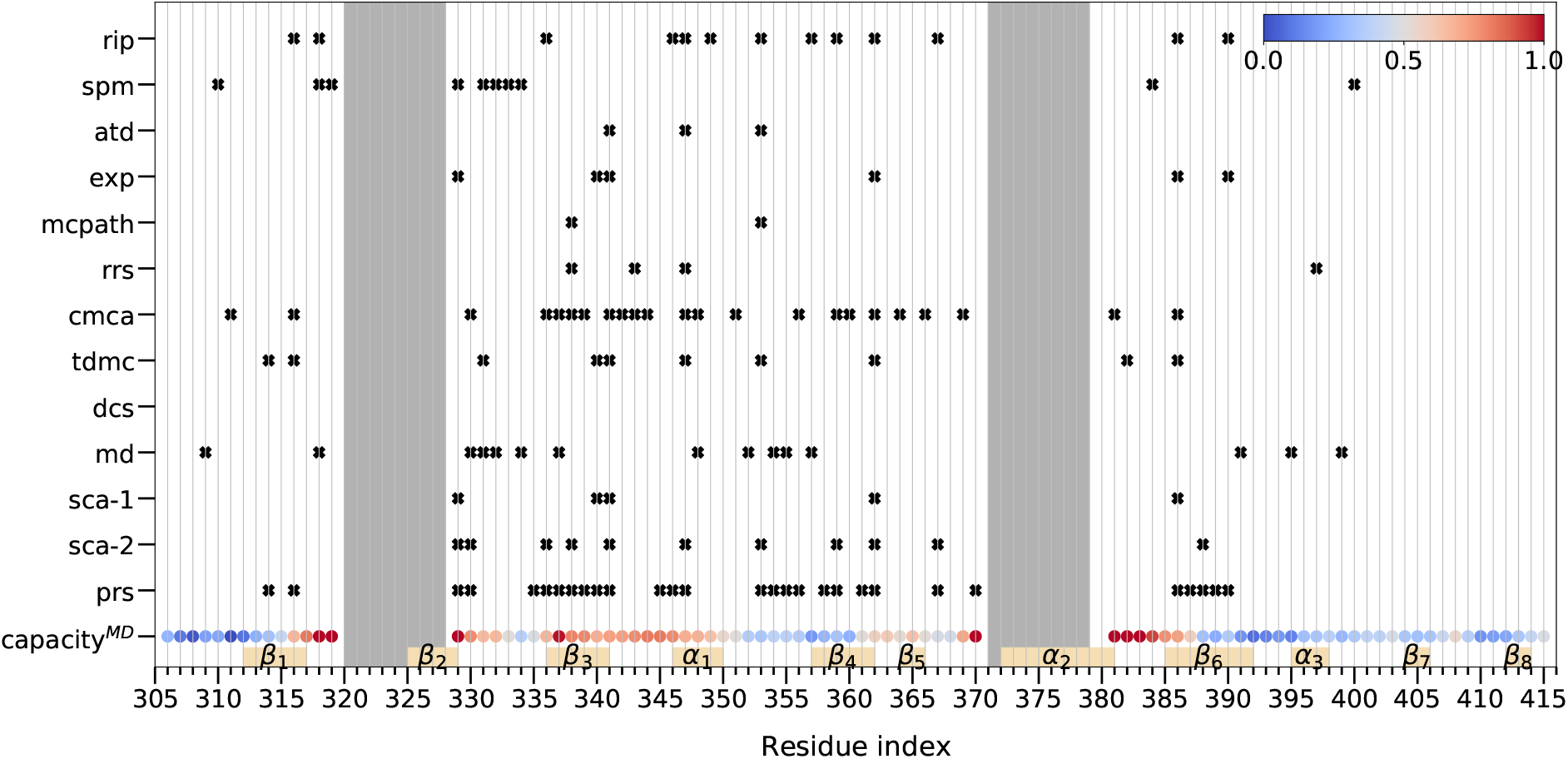
Capacity values from BP excitation and the critical residues identified by various methods. The gray areas indicate the residues on which the external force is applied in the capacity analysis. Capacity^MD^ refers to the results obtained with ***H***^MD^. Abbreviations: prs - perturbation response scanning [45], exp - experimental [46], sca-1 - statistical coupling analysis [47], sca-2 - statistical coupling analysis [48], atd - anisotropic thermal diffusion [49], spm - structural perturbation method [50], rip - rotamerically induced perturbation [51], md - molecular dynamics [42], dcs - deep coupling scan [52], tdmc - thermodynamic double mutant cycle [53], cmca - conservation mutation correlation analysis [54], rrs - rigid-residue scan [55], mcpath - Monte Carlo path [56].

In order to investigate role of frequency in determining the allosteric responses, we present the frequency dependent SNR profiles in Figure 10(a). The side views of the figure are in Figure S6a. The frequency dependence of the SNRs for the individual residues are also shown in Figure S7. Although the majority of the residues display monotonic decrease in SNR with increasing frequency, it is worth noting that some of the residues exhibit a peak, a sort of *resonance*, around a certain frequency value, that is specific to the residue. The residues with distinct frequency response are presented in Figure 10(b). The responses of the residues differ in terms of the magnitude of SNR, the slope of the decay with frequency, and the existence and frequency location of a resonance at higher frequencies. For instance, GLY 329, which is one of the few consistently identified critical allosteric residues in Figure 9 has a relatively high response until the high end of the spectra. The resonance around 2 THz is also worth noting for GLY 329. The resonance at about the same frequency is significantly more pronounced for GLN 384. The other listed residues display resonances with smaller magnitudes at high frequencies. With these results, we underline that the frequency decomposed SNR analysis provides a more detailed picture of allostery, paving the way for gaining a better understanding of allosteric mechanisms.

**Figure 10:**
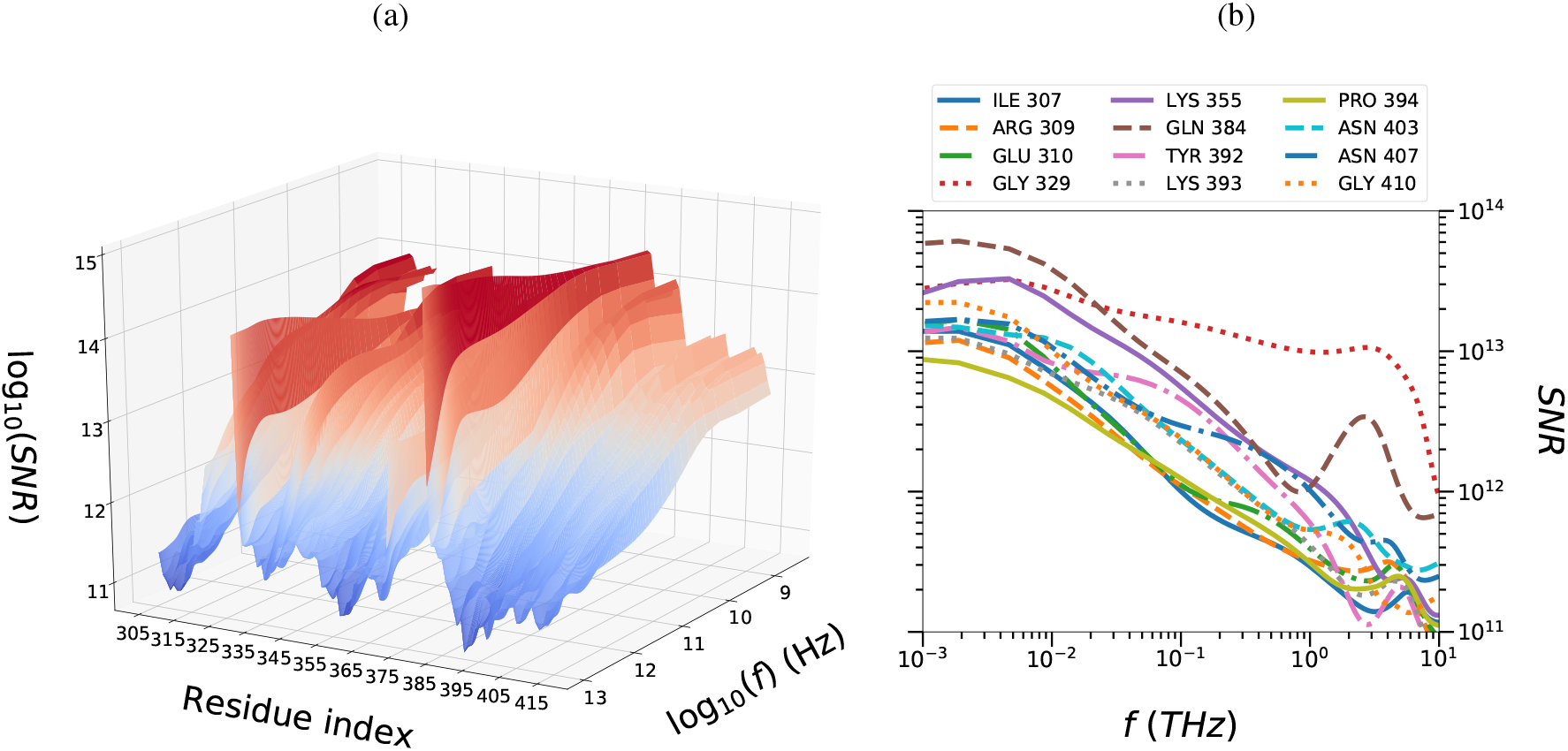
(a) Signal-to-Noise ratio (SNR) with ***BP*** excitation using ***H***^MD^, (b) Selected residues with characteristic frequency response.

## 5 Conclusion

The aim of the research presented in this paper was to investigate the phenomenon of allostery using well established tools of telecommunication systems, namely frequency decomposed SNR and channel capacity based analyses. To that end, we analyzed the displacements due to both noise forces and external excitations—capturing the effect of the ligand—separately. We proposed two related analysis schemes, termed as *per residue scan* and *binding pocket excitation*, in order to identify the residue pairs that are likely to interact allosterically, and the residues that are affected in a significant manner by a particular ligand-binding event, respectively. The frequency domain representations employed by the proposed methods lead to an alternate view into allostery, by emphasizing the effect of perturbation frequency in the SNR response. We have shown that the response of some of the residues exhibit a resonance at specific, characteristic frequencies. Thus, a full spectral analysis of the responses to perturbations leads to the speculative but potentially significant conclusion that the key mechanism underlying allostery is robust signal transmission despite noise at specific frequencies. The frequency-decomposed approach further allows the analysis of equilibrium fluctuations in the absence of the ligand. We proposed an alternative technique to compute mean square fluctuations in a frequency-decomposed manner. In all of our computations, we adopted a Langevin formulation and considered different options to obtain the second-order partial derivatives of the potential energy function, namely the Hessian. The equilibrium analysis suggests that although the mean square fluctuation profiles obtained from molecular dynamics simulations and elastic network models are quite similar, their frequency distributions have rather distinct characteristics. Therefore, the results from simplified models should be interpreted with caution.

To the best of our knowledge, we presented the first comprehensive application of SNR and channel capacity based analysis methods in the context of allostery. However, the full potential of the proposed methods has not been fully explored yet. More detailed analyses are required to determine the full benefits and limitations. Their usage and interpretations were demonstrated for the well-studied PDZ3 protein. It is unfortunate that there is no consensus in the literature on the residues that play a significant role in the allosteric behavior of this protein. However, reaching a consensus seems to be exceptionally difficult when the tremendous challenges involved in both experimental and computational techniques aimed at deciphering allostery are considered.

## Data and Code Availability

The proposed methods were implemented in Python and available at https://github.com/yabozkurt/protein_cap. ProDy package [57] is utilized for parsing the PDB files and for the construction of the ANM model, and MDAnalysis is used for structure alignments [58].

## Acknowledgments

YBV and AD would like to thank Prof. Burak Erman (Koc University) for various discussions. YBV acknowledges financial support from the Scientific and Technological Research Council of Turkey (TUBITAK) under the 2211 graduate scholarship program. A part of the paper was written while YBV was a visiting student at the Max Planck Institute for Polymer Research, financially supported by the TUBITAK 2214-A program on international collaborative research.

## Supporting Information for

### S.1 SNR and Channel Capacity

**Figure S1:**
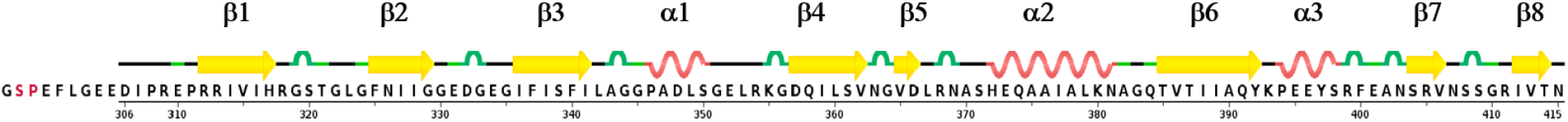
The secondary structure assignments from DSSP (definition of secondary structure of proteins) [59] of the holo form of the PDZ3 protein (PDBID: 1be9). Image is from the RCSB PDB (www.rcsb.org) [60].

**Table S1:**
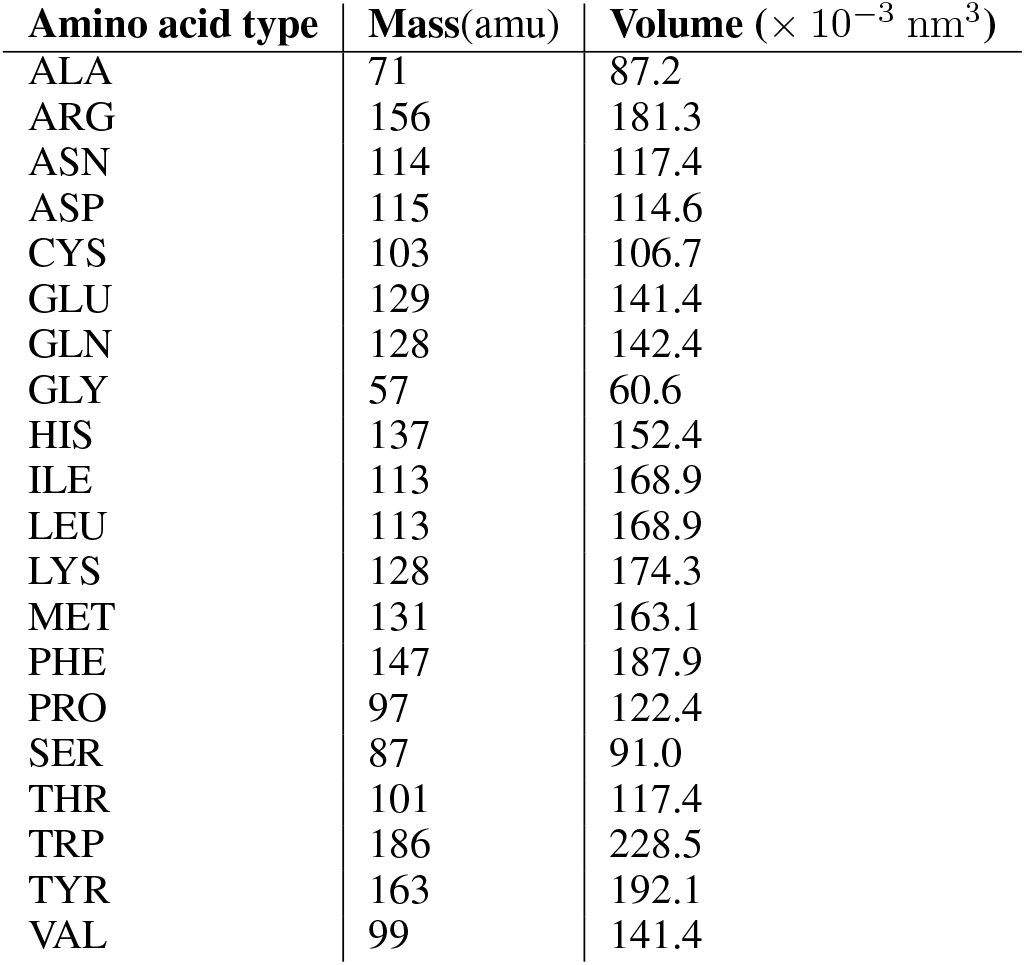
Mass and volume values of the amino acids

| Amino acid type | Mass(amu) | Volume ($\times 10^{-3} \text{ nm}^3$) |
| --- | --- | --- |
| ALA | 71 | 87.2 |
| ARG | 156 | 181.3 |
| ASN | 114 | 117.4 |
| ASP | 115 | 114.6 |
| CYS | 103 | 106.7 |
| GLU | 129 | 141.4 |
| GLN | 128 | 142.4 |
| GLY | 57 | 60.6 |
| HIS | 137 | 152.4 |
| ILE | 113 | 168.9 |
| LEU | 113 | 168.9 |
| LYS | 128 | 174.3 |
| MET | 131 | 163.1 |
| PHE | 147 | 187.9 |
| PRO | 97 | 122.4 |
| SER | 87 | 91.0 |
| THR | 101 | 117.4 |
| TRP | 186 | 228.5 |
| TYR | 163 | 192.1 |
| VAL | 99 | 141.4 |

Table S2 summarizes the critical residues identified in previous work using a wide range of computational methods as well as experimental techniques for the PDZ3 domain protein PSD-95. The summaries provided in [43] and [45] were utilized in the preparation of the table.

**Figure S2:**
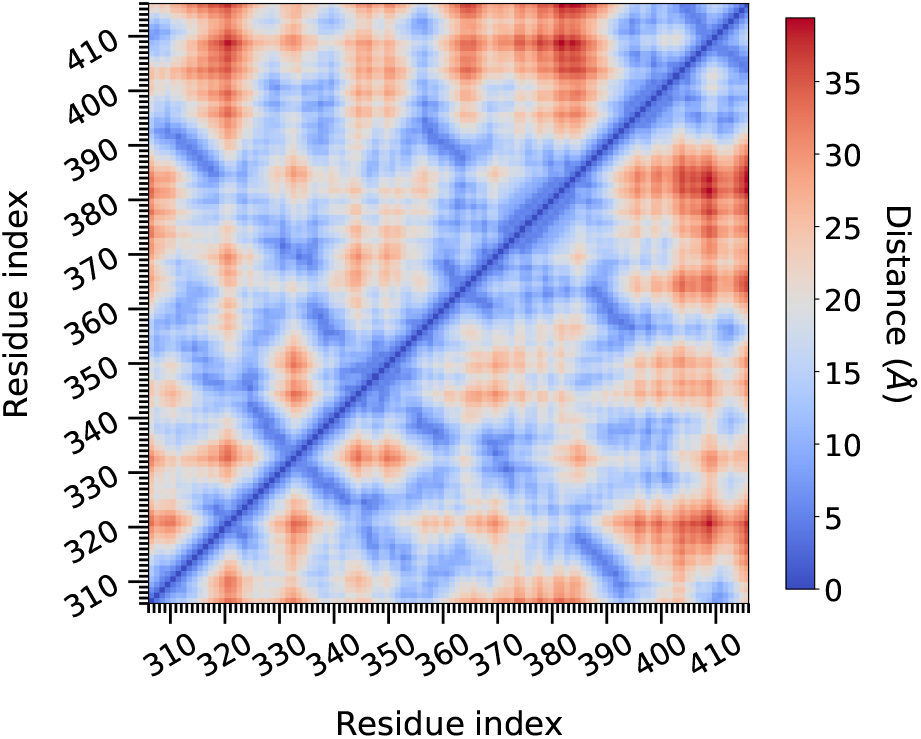
Pairwise distances between the residues

**Table S2:** Critical residues identified by various methods

| Method | Critical residues |
| --- | --- |
| Perturbation response scanning [45] | 314, 316, 326, 327, 328, 329, 330, 335, 336, 337, 338, 339, 340, 341, 345, 346, 347, 353, 354, 355, 356, 358, 359, 361, 362, 367, 370, 372, 375, 379, 386, 387, 388, 389, 390 |
| Experimental [46] | 325, 328, 329, 340, 341, 362, 372, 376, 380, 386, 390 |
| Statistical coupling analysis. 1 [47] | 322, 325, 329, 340, 341, 362, 372, 376, 380, 386 |
| Statistical coupling analysis. 2 [48] | 323, 324, 325, 327, 328, 329, 330, 336, 338, 341, 347, 353, 359, 362, 367, 372, 375, 376, 379, 388 |
| Anisotropic thermal diffusion [49] | 325, 327, 341, 347, 353, 372 |
| Structural perturbation method [50] | 310, 318, 319, 320, 323, 327, 329, 331, 332, 333, 334, 372, 376, 380, 384, 400 |
| Rotamerically induced perturbation [51] | 316, 318, 323, 325, 336, 346, 347, 349, 353, 357, 359, 362, 367, 375, 378, 379, 386, 390 |
| MD [42] | 309, 318, 322, 323, 324, 326, 327, 328, 330, 331, 332, 334, 337, 348, 352, 354, 355, 357, 373, 380, 391, 395, 399 |
| Deep coupling scan [52] | 372, 375, 376, 379 |
| Thermodynamic double mutant cycle [53] | 314, 316, 323, 325, 327, 328, 331, 340, 341, 347, 353, 362, 372, 375, 376, 377, 378, 380, 382, 386 |
| Conservation mutation correlation analysis [54] | 311, 316, 326, 327, 330, 336, 337, 338, 339, 341, 342, 343, 344, 347, 348, 351, 356, 359, 360, 362, 364, 366, 369, 376, 378, 381, 386 |
| Rigid-residue scan [55] | 338, 343, 347, 397 |
| Monte Carlo path [56] | 325, 327, 338, 353, 372 |

**Figure S3:**
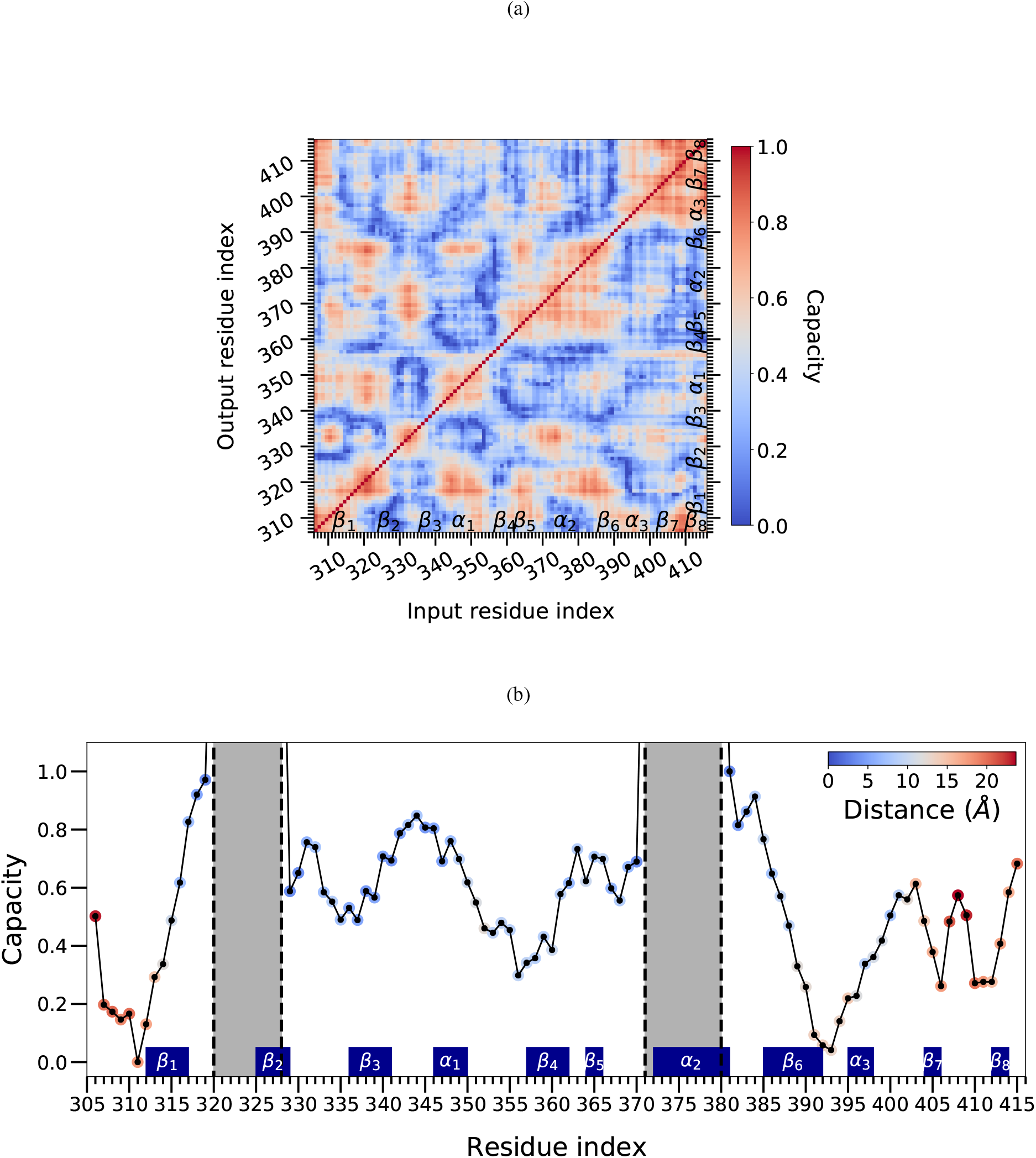
(a) Per residue scan and (b) BP excitation with ***H***^ANM^

**Figure S4:**
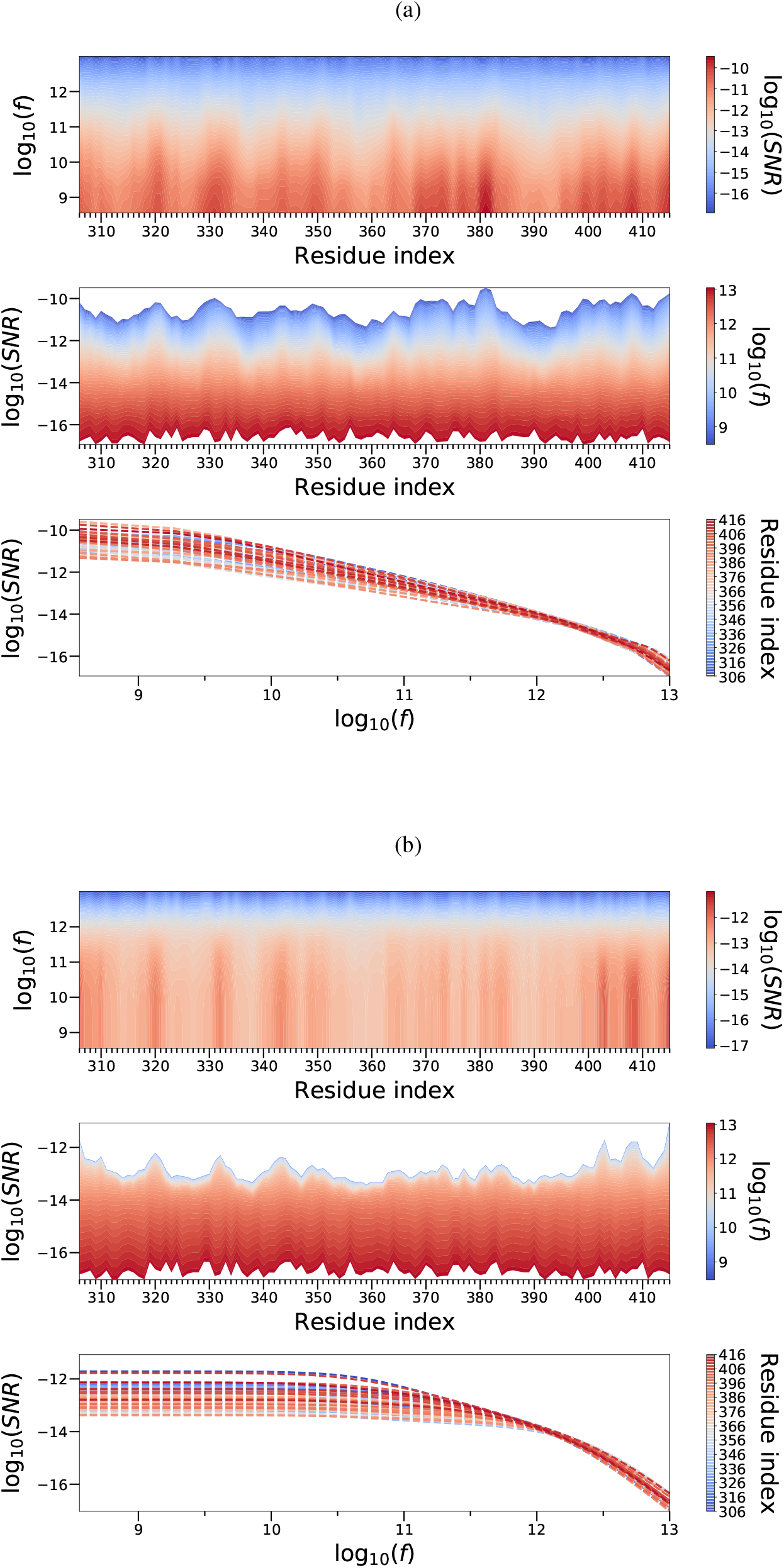
PSD profile for the equilibrium fluctuations (a) ***H***^MD^ (b) ***H***^ANM^

**Figure S5:**
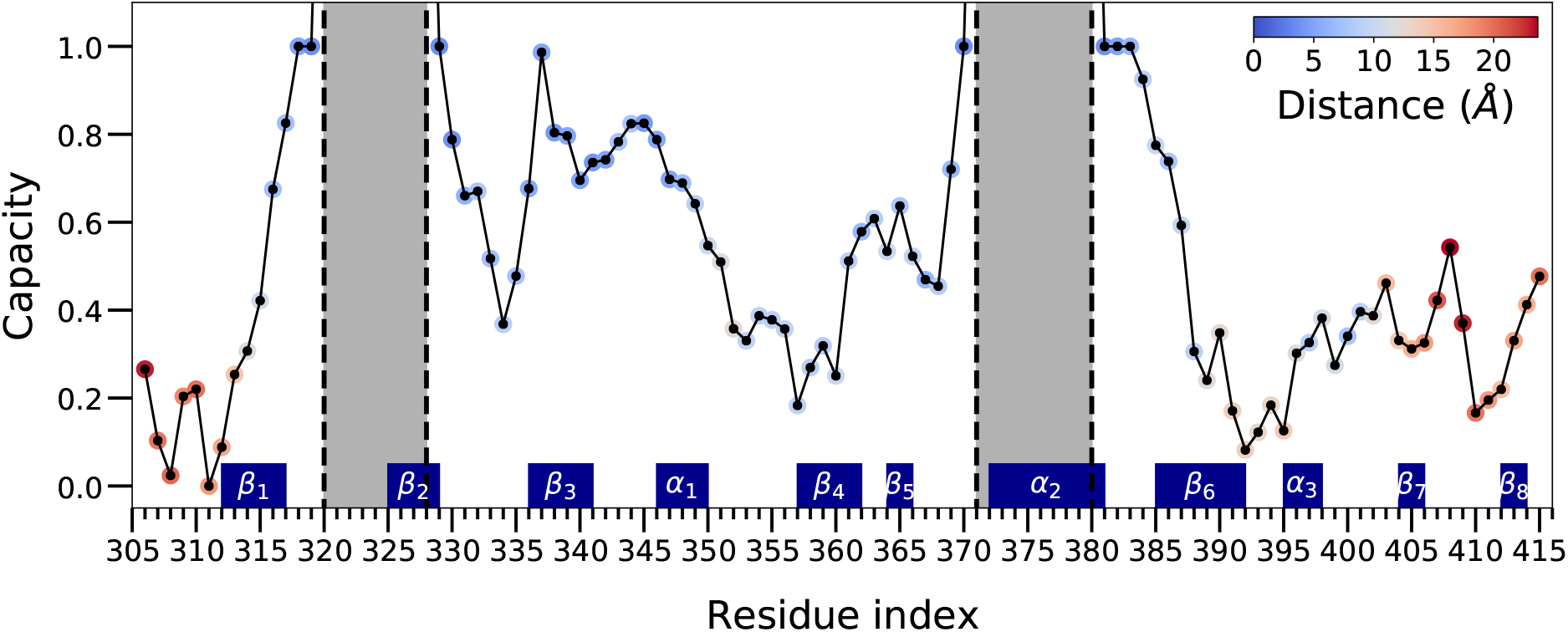
Normalized capacity values in BP excitation mode using ***H***^MD^. The colorbar shows the minimum distance between a residue and one of the BP residues.

**Figure S6:**
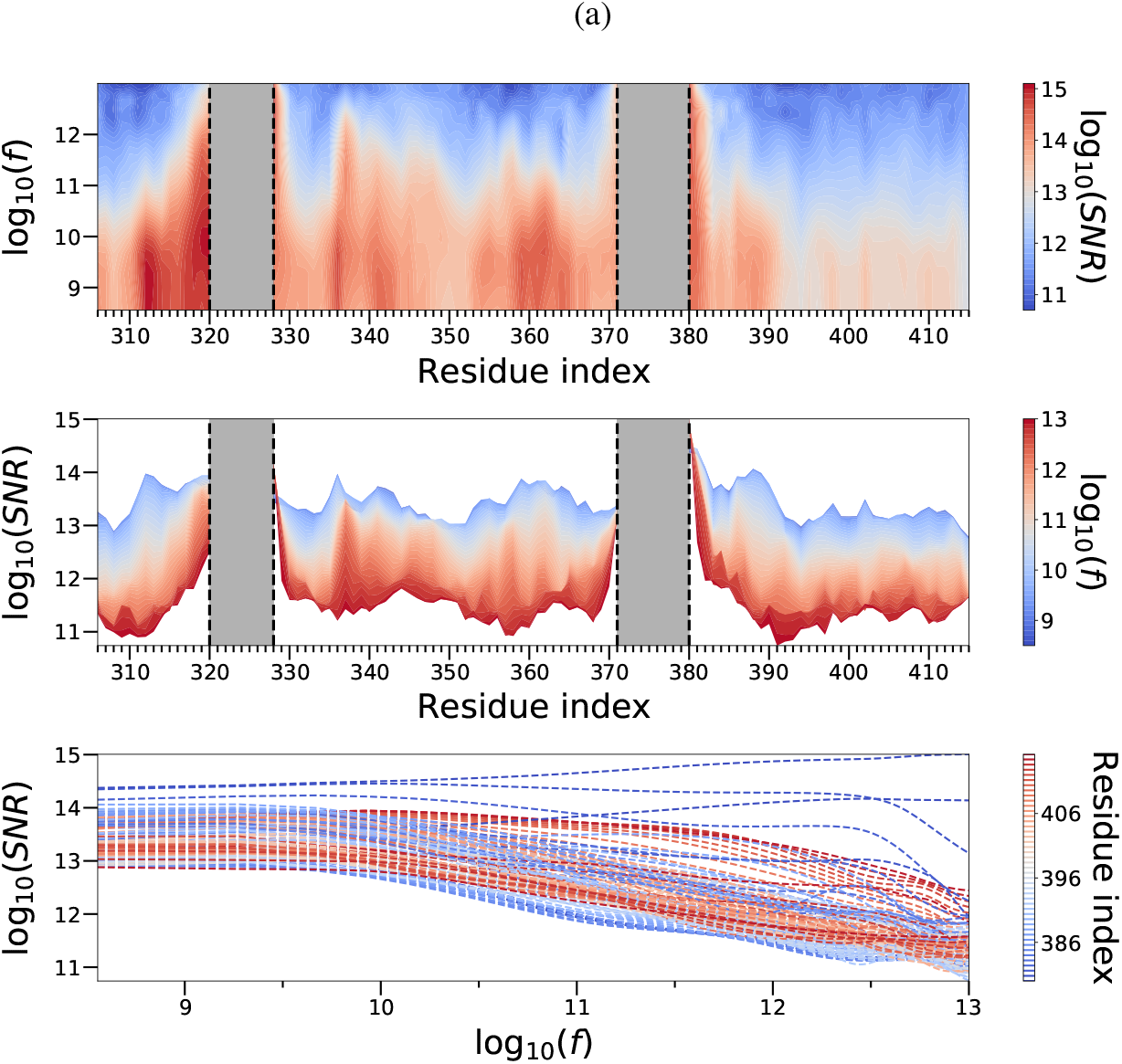
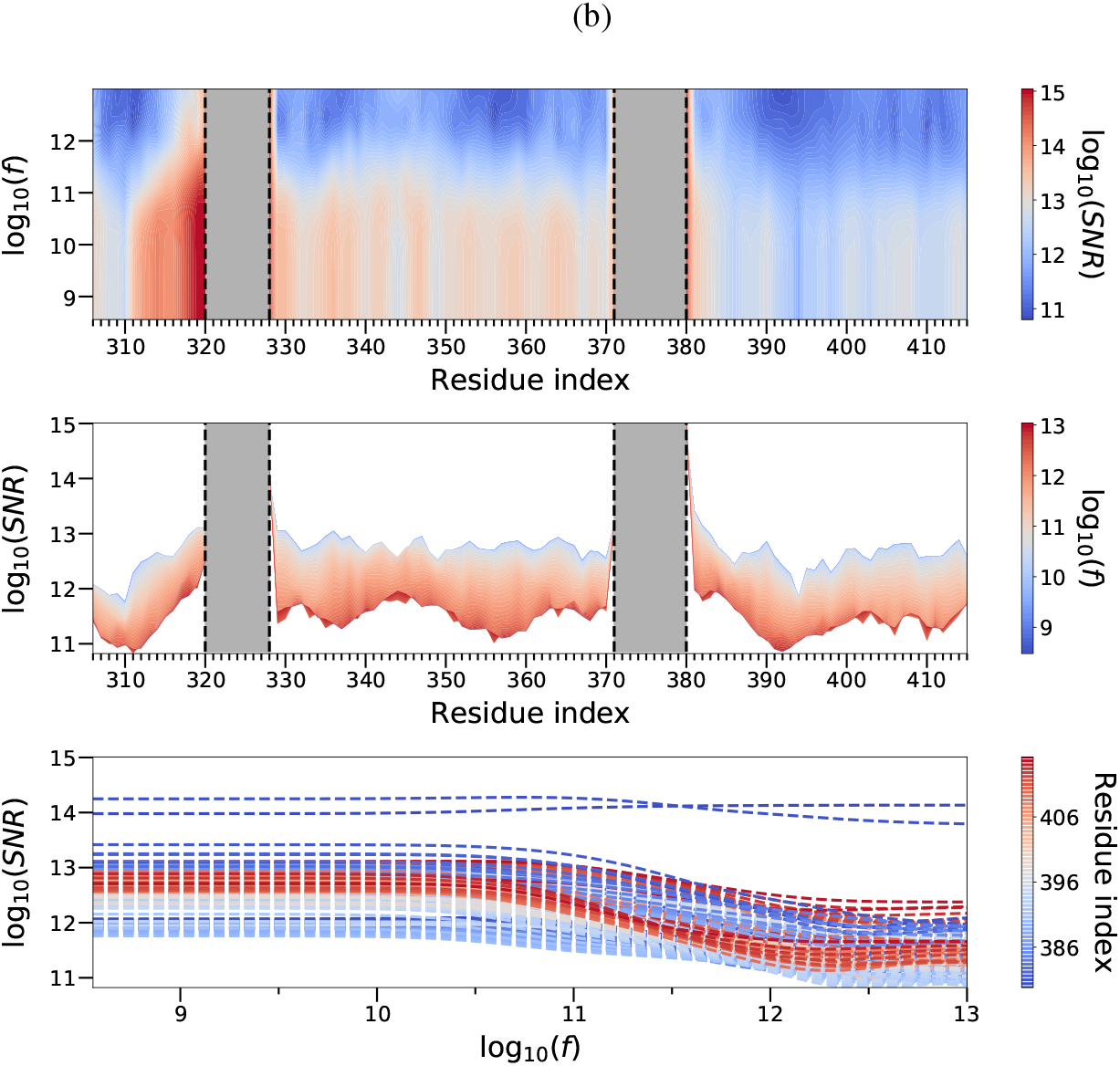
Binding pocket excitation with (a) ***H***^MD^ (b) ***H***^ANM^.

**Figure S7:**
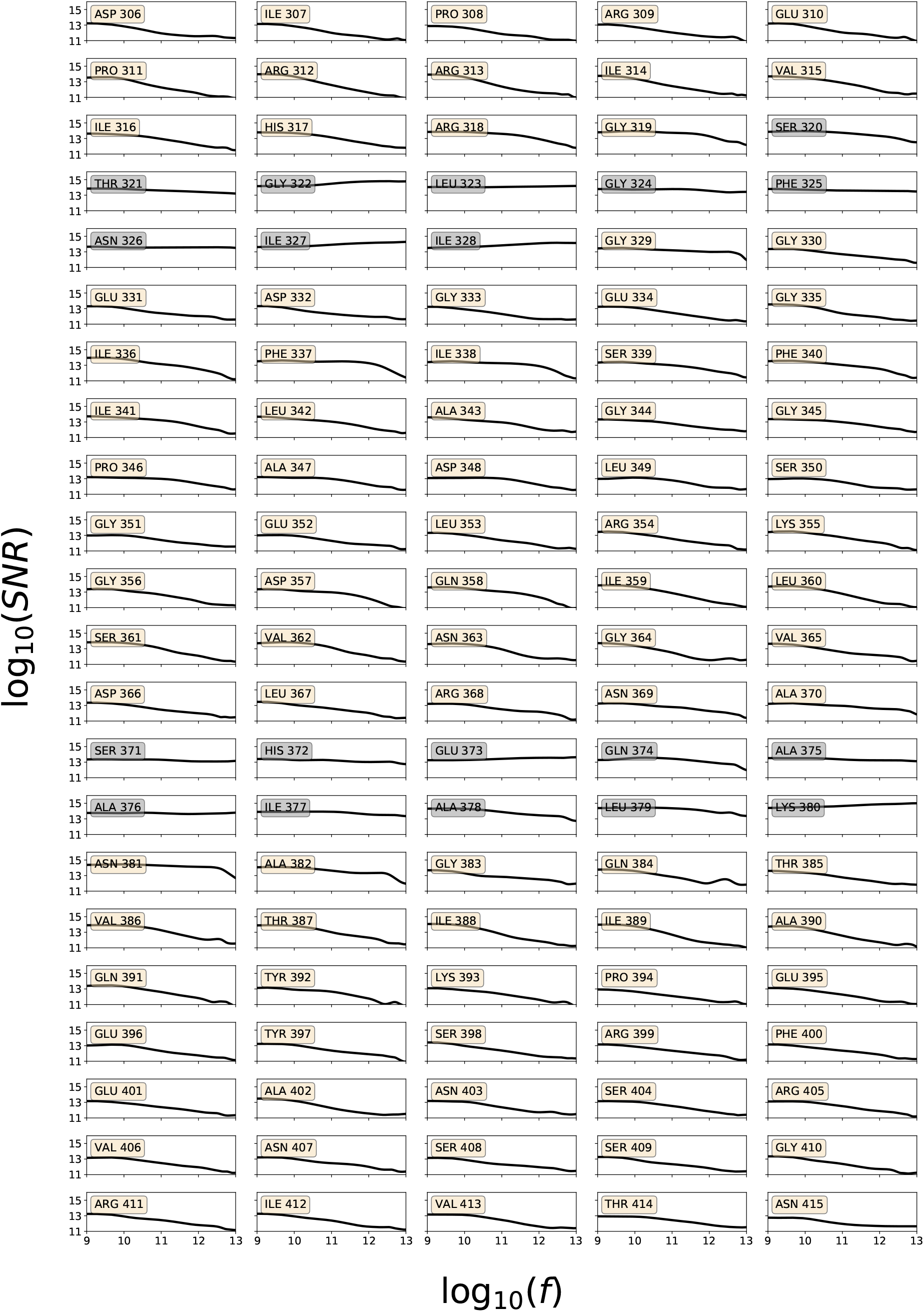
Individual frequency response of the residues in BP excitation with ***H***^MD^.

### S.2 Constrained Least Squares

The derivation of the solution for the following optimization problem described in Equation 12 is adopted from [61]

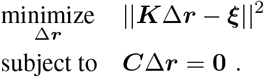

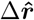 is a solution.

Lagrangian function with Lagrange multipliers *λ*_1_, *λ*_2_, …, *λ*_6_:

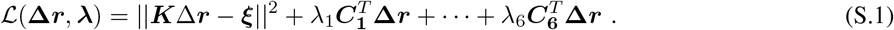

Two optimality conditions are:

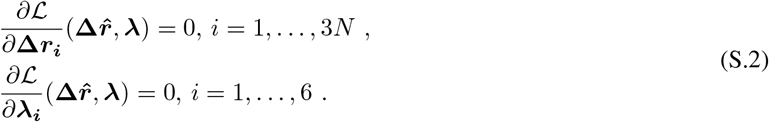

The second optimality condition yields: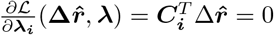.

The first set of conditions yields:

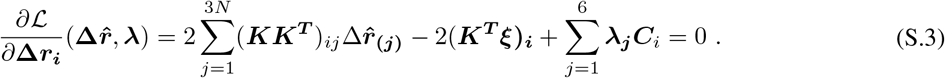

In matrix-vector form

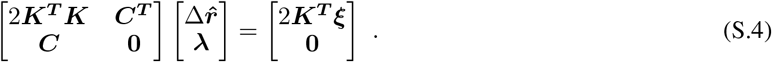

### S.3 Lyapunov Formulation

This section presents a Lyapunov-equation based approach in order to show that the mean square fluctuations are independent of the values of the friction coefficients. The rationale behind this discussion is that random noise forces and the viscous friction forces have counteracting effects, as dictated by the Fluctuation-Dissipation theorem [17, 18].

Equation 4 can be recast as 6*N* coupled first order differential equations with the state space vector ***v***

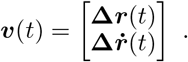

Under no external force, Equation 4 can be expressed as an initial value problem, with initial conditions ***v***(0), as follows

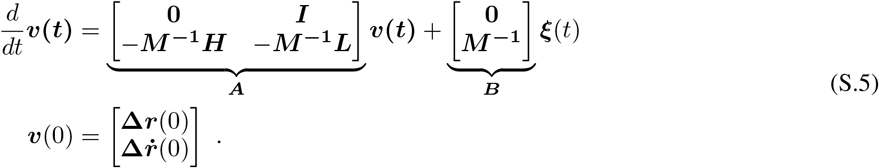

Due to the fact that white noise ***ξ***(*t*) can be expressed as the formal time derivative of a Wiener Process, ***W***, Equation S.5 is written in differential form as follows

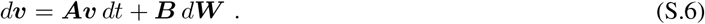

Noise characteristics are as follows:

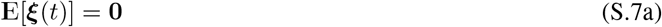

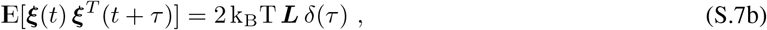

where E[·] denotes the probabilistic expectation operator. Taking the expectation of both sides of Equation S.6, and using the property in Equation S.7a

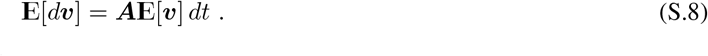

The zero time-lag correlation matrix is defined as

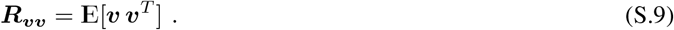

From Equations S.6 and S.9, and using Ito’s product rule [62] (from stochastic Ito calculus), the following equation is derived

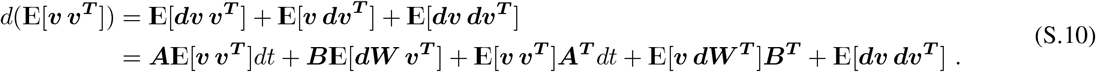

The second and fourth terms in Equation S.10 are zero due to the fact that ***v*** and ***dW*** are uncorrelated [63], i.e.,

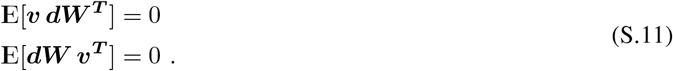

The addditional term in the stochastic calculus product rule in Equation S.10 can be expanded as

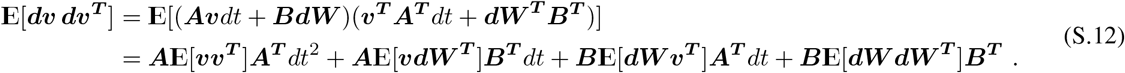

The first three terms in Equation S.12 are set to zero due to the following that follow from rules of stochastic Ito calculus (The second and third terms vanish, also due to Equation S.11.)

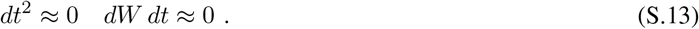

From the noise properties in Equation S.7b

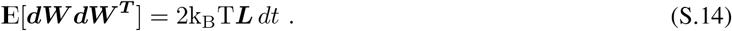

Then, Equation S.10 becomes

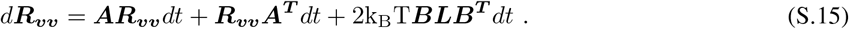

Zero time-lag covariance matrix ***C***_***vv***_ is defined as

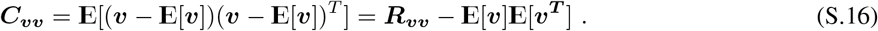

Differential form of ***C***_***vv***_:

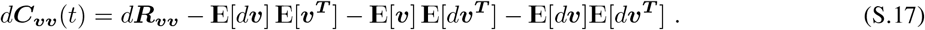

Utilizing Equation S.8 and expanding *d****R***_***vv***_ according to Equation S.15 yields

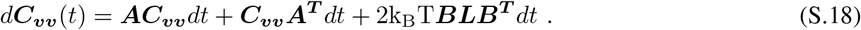

In the limit as time goes to infinity, ***v***(*t*) approaches to a (wide-sense) stationary process where E[***v***] and ***C***_***vv***_ become independent of time and we obtain Equation S.19b, a *Lyapunov equation* [64].

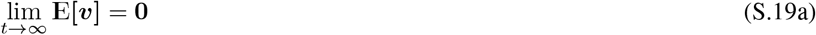

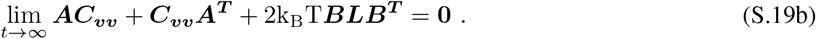

We note that, due to Equation S.19a, ***C***_***vv***_ = ***R***_***vv***_, however we stick to the covariance notation. Using the symmetricity of covariance matrices, ***C***_***vv***_ is structured in block form as

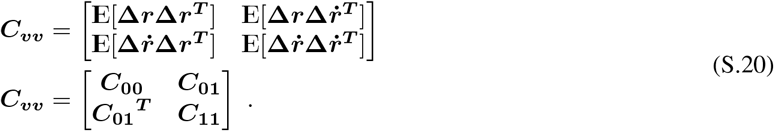

Inserting ***A*** defined in Equation S.5 and the block form of ***C***_***vv***_, with ***H*** = ***H***^***T***^, Equation S.19b expands as

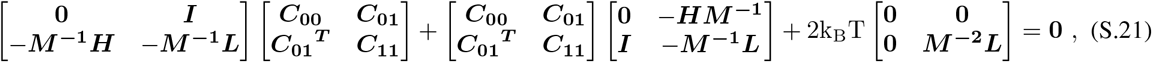

which can be further manipulated as

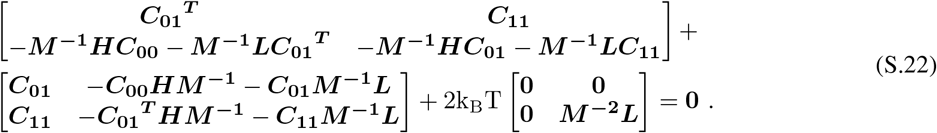

Equation S.22 leads to the following set of equations

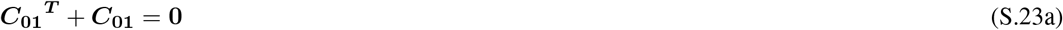

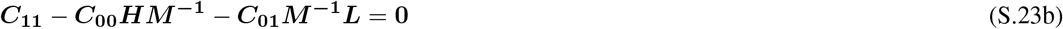

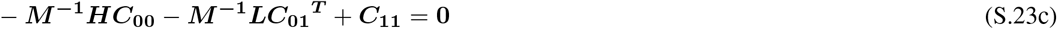

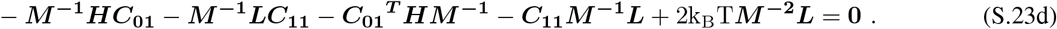

Equation S.23a implies that *C*_01_ = −***C***_**01**_^***T***^, i.e., ***C***_**01**_ is antisymmetric. Then, the above set of equations yield

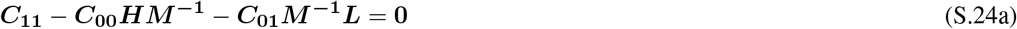

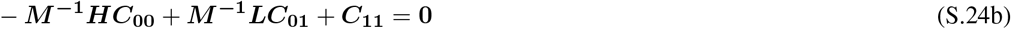

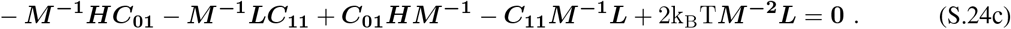

From Equations S.24a and S.24b

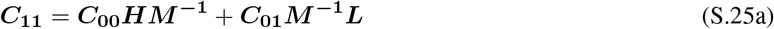

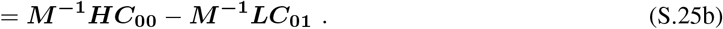

Noting the symmetricity of covariance matrices: ***C***_**00**_ = ***C***_**00**_^***T***^ and ***C***_**11**_ = ***C***_**11**_^***T***^. Then, inserting Equation S.25 into Equation S.24c gives

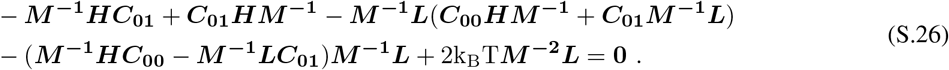

Multiplying the above by ***M*** both from the left and right results in

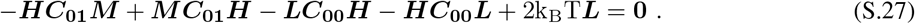

When ***C***_00_ = k_B_T***H***^**−1**^, Equations S.25a and S.25b together read

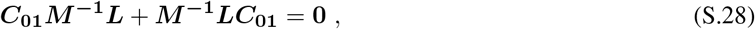

implying that *C*_01_ = 0. We then observe that ***C***_00_ = k_B_T***H***^**−1**^ satisfies Equation S.27, based on the fact that the first two terms are equal to zero with this form of solution for ***C***_00_, as shown above. We also note that ***M***, ***M*** ^−1^, ***L*** are all diagonal matrices with positive-valued entries. This shows that the mean square displacements are independent of the chosen ***L***. We note also that ***H*** is singular, thereby its inverse is calculated either as a pseudo-inverse, or alternatively, by adapting the constraint-based solution that is introduced in Section S.2. The frequency-dependent terms are zero in the transfer function (***T***) definition at zero frequency in Equation 9, i.e., ***T* (0)** = ***H***^**−1**^. To address the singularity problem, rotational and translational constraints are introduced into the transfer function, leading to Equation 14. Then, the constrained transfer function at zero frequency, ***T***_***c***_**(0)**, denoted as ***H***_***c***_^**−1**^, can be calculated as

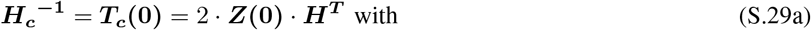

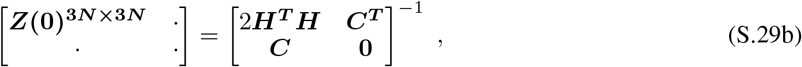

where ***C*** corresponds to the constraints matrix, defined in Equation 11.

As an alternative approach in order to demonstrate that the mean square displacements are independent of the friction values, Figure S8 shows the numerical solution of Equation S.19b for ***C***_00_. Due to the previously mentioned invertibility issues of ***H***, Lyapunov equation does not have a unique solution with matrix ***A*** arising from a rank-deficient ***H***. In order to overcome this problem in the numerical solution, ***H***_***c***_ is utilized instead. Figure S8 is drawn for various ***L*** values. The same figure is obtained in all trials. Therefore, it can be concluded that due to the link between thermal noise and viscous friction, residue fluctuations are independent of the friction values.

**Figure S8:**
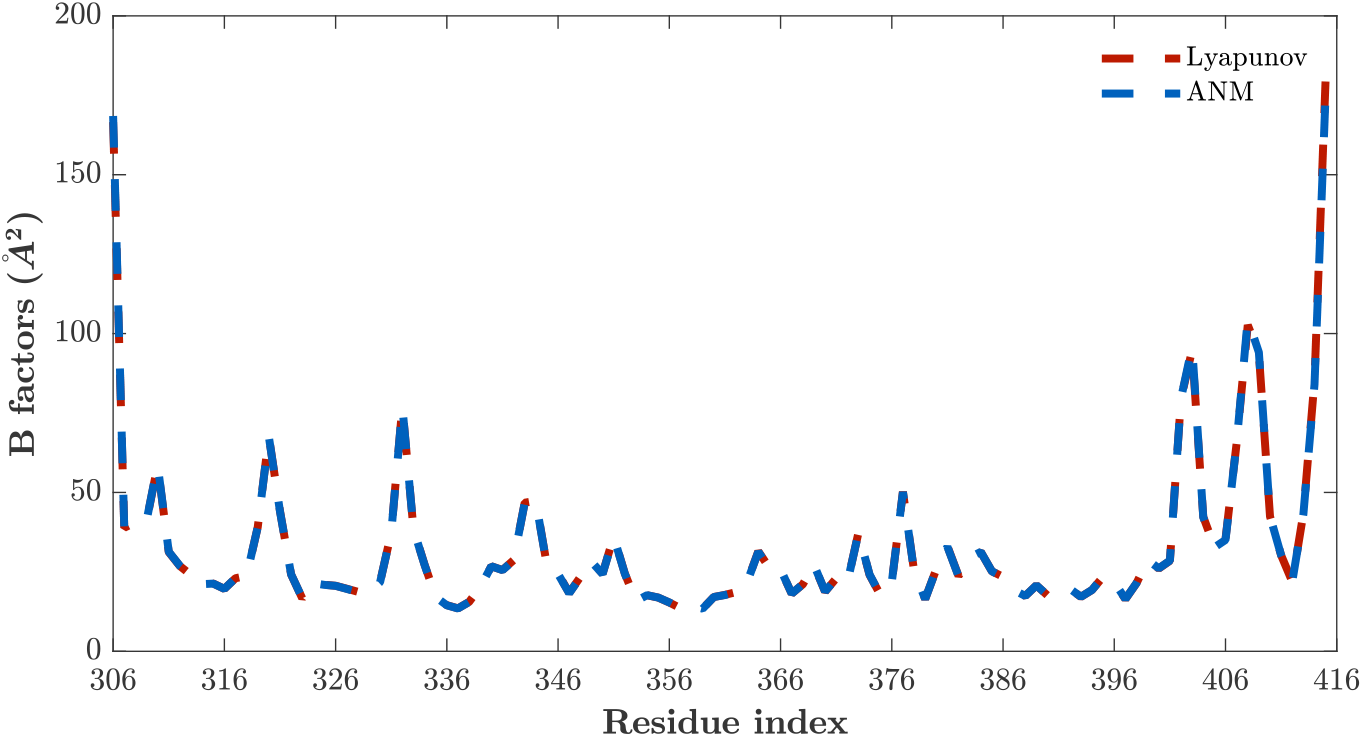
Numerical Solution of Lyapunov equation

